# ZFAND6 is a subunit of a TRAF2-cIAP E3 ubiquitin ligase complex essential for mitophagy

**DOI:** 10.1101/2024.03.27.586763

**Authors:** Kashif Shaikh, Melissa Bowman, Sarah M. McCormick, Linlin Gao, Jiawen Zhang, John Tawil, Arun Kapoor, Ravit Arav-Boger, Young Bong Choi, Andrew Pekosz, Sabra L. Klein, Matthew Lanza, Julie C. Fanburg-Smith, Adolfo García-Sastre, Christopher C. Norbury, Zissis C. Chroneos, Edward W. Harhaj

## Abstract

The A20 ubiquitin-editing enzyme is a critical negative regulator of NF-κB signaling and inflammation. While the mechanisms by which A20 restricts inflammation have been extensively studied, the physiological functions of other A20-like proteins are largely unknown. Here, we report a previously unknown function of the A20 family member ZFAND6 as a novel regulator of mitophagy. Deletion of ZFAND6 in bone marrow-derived macrophages (BMDMs) promotes the upregulation of reactive oxygen species (ROS) and the accumulation of damaged mitochondria due to impaired mitophagy. Consequently, mitochondrial DNA (mtDNA) is released into the cytoplasm, triggering the spontaneous expression of interferon-stimulated genes (ISGs) in a cGAS-STING dependent manner, which leads to enhanced viral resistance *in vitro*. However, mice lacking ZFAND6 exhibit increased morbidity and mortality upon challenge with a sublethal dose of influenza A virus (IAV) due to impaired myeloid cell activation and diminished type I IFN signaling. Mechanistically, ZFAND6 bridges a TRAF2-cIAP1 interaction, which is required for the initiation of ubiquitin-dependent mitophagy. Our results suggest that ZFAND6 is a subunit of a TRAF2-cIAP E3 ligase complex that promotes the clearance of damaged mitochondria by mitophagy to maintain mitochondrial homeostasis.

## Main

Activation of pattern recognition receptors (PRRs) is crucial for inflammatory responses to control microbial infections. However, aberrant activation of PRRs leads to chronic inflammation and is associated with autoimmunity^1, 2^ and neurological disorders^3, 4^. To counteract the deleterious effects of inflammation, mammalian organisms have evolved complex mechanisms to tightly regulate inflammatory responses. However, in the settings of autoimmune or autoinflammatory disease, these pathways can become perturbed, resulting in chronic inflammatory signaling. A20 (also known as TNFAIP3) is a ubiquitin-editing enzyme and a key negative regulator of NF-κB signaling^5, 6,7^. A20 contains an Ovarian Tumor (OTU) domain^8^ and seven C_2_/C_2_ zinc finger (ZF) domains^9^. A20 inhibits TNF-induced NF-κB signaling by removing Lys^63^(K63)-polyubiquitin chains from RIPK1 using its OTU domain, followed by ZF4-dependent conjugation with Lys^48^(K48)-polyubiquitin chains to trigger RIPK1 degradation^7^. A20 is well established as a critical regulator of inflammatory and cell death pathways that is essential to prevent unrestrained inflammation and autoimmunity^10^. In addition to A20, there are nine other A20 family members with A20-like ZF domains encoded in the human genome; however, their physiological functions are largely unknown^11^.

ZFAND6 (also known as AWP1) is a member of the A20 family that contains an A20-like ZF domain as well as an AN1 ZF domain^11^. ZFAND6 was first identified as a binding partner of the protein kinase C related kinase 1 (PRK1)^12^. *Zfand6* mRNA was also shown to be induced in adherent primary monocytes and was proposed to function as an inhibitor of NF-1B^13^. ZFAND6 can interact with the E3 ubiquitin ligase TRAF2 and regulate tumor necrosis factor (TNF)-induced NF-κB signaling through its A20-like ZF domain^14^. Furthermore, ZFAND6 has been shown to mitigate the generation of reactive oxygen species (ROS) in breast cancer cells^15^. In plants, proteins containing both A20-like and AN1 ZF domains are involved in the stress response^16^. In humans, genome-wide association studies (GWAS) have implicated the *Zfand6* gene as a potential susceptibility locus for type 2 diabetes^17^. However, the functions and physiological roles of ZFAND6 are unknown.

Mitochondrial autophagy (mitophagy) is a process whereby damaged mitochondria are removed through lysosomal degradation^18^. In metabolically active cells such as macrophages, mitochondria become damaged due to oxidative stress, which necessitates their clearance through mitophagy. Impairments in mitophagy could lead to the accumulation of damaged mitochondria and ROS^19^, which results in the release of mitochondrial damaged-associated molecular patterns (DAMPs), leading to aberrant activation of PRRs^20, 21^. Mitochondria undergo mitophagy either by receptor-mediated mitophagy or ubiquitin-mediated mitophagy^18^. In receptor-mediated mitophagy, mitophagy receptors such as BNIP3 and NIX accumulate on damaged mitochondria and recruit autophagosomes through binding to ATG8 family members to degrade mitochondria in lysosomes^22^. Ubiquitin-mediated mitophagy is classically driven by the PINK-Parkin pathway^18^. PINK1, a serine-threonine kinase, is recruited to damaged mitochondria and is stabilized on the outer mitochondrial membrane (OMM) by the loss of mitochondrial membrane potential^23^. PINK1 then recruits and phosphorylates the E3 ubiquitin ligase Parkin^24^, and also phosphorylates ubiquitin (Ub) on Ser65 to promote Parkin activation^25, 26^. This leads to the recruitment of selective autophagy receptors such as NDP52, Optineurin and p62/SQSTM1 to the damaged mitochondria, which aids in their recruitment to the mitophagosomes for their fusion with the lysosome for proteolytic degradation^27^.

TNF receptor-associated factor 2 (TRAF2), a member of the TRAF (TNF receptor associated factors) family of proteins, mediates NF-κB signaling as a component of TNFR signaling complexes to promote the activation of inhibitor of IκB kinase (IKK)^28^. Furthermore, TRAF2, together with TRAF3 and cellular inhibitor of apoptosis 1 and 2 (cIAP1, 2) proteins, constitute an E3 ligase complex that regulates non-canonical NF-κB signaling^29^. TRAF2 has recently emerged as a key regulator of mitophagy although the underlying mechanisms are unclear^30^. TRAF2 has been proposed to mediate Parkin-independent ubiquitin-mediated mitophagy in cardiac myocytes^31, 32^. However, whether TRAF2 functions independently or as part of an E3 ligase complex in this function is unknown. In this manuscript, we identify ZFAND6 as a novel regulator of mitophagy. ZFAND6 functions as an adaptor protein to facilitate cIAP1-TRAF2 interactions upon mitophagy induction. Loss of ZFAND6 leads to the accumulation of damaged mitochondria which release DAMPs that activate the cGAS-STING pathway.

## Results

### Loss of ZFAND6 triggers the upregulation of ISGs

Mouse *Zfand6* consists of 7 exons, of which exons 2-7 encode the ZFAND6 protein. To understand the physiological functions of ZFAND6, we generated mice lacking *Zfand6* by deleting and replacing exons 2-4 with a neomycin (Neo) cassette (Extended Data Fig. 1A-B). *Zfand6*^−/−^ mice appeared normal and did not exhibit any obvious developmental or immune defects (data not shown). An in-depth characterization of the immune system of *Zfand6*^−/−^ mice will be described elsewhere. We generated WT and *Zfand6*^−/−^ murine embryonic fibroblasts (MEFs) and bone marrow-derived macrophages (BMDMs) and confirmed that ZFAND6 expression was absent in knockout cells (Extended Data Fig. 1C-D). To determine if ZFAND6 regulated Toll-like receptor 4 (TLR4) or RIG-I-like receptor (RLR) signaling we harvested BMDMs from wild-type and *Zfand6*^−/−^ mice and stimulated with either bacterial lipopolysaccharide (LPS) or infected with influenza A virus (IAV) and performed bulk RNA-Seq analysis (Fig. 1A). Gene-set enrichment analysis revealed the spontaneous expression of over 100 interferon-stimulated genes (ISGs) in *Zfand6*^−/−^ BMDMs, and these genes were also upregulated in knockout BMDMs after LPS stimulation or IAV infection (Fig. 1B-C). The increased expression of select ISGs (*Ifit3*, *Isg15*, *Trim34b*, and *Mx1*) in ZFAND6-deficient cells was validated by real-time qRT-PCR in MEFs and BMDMs (Fig. 1D-E).

**Figure 1.**
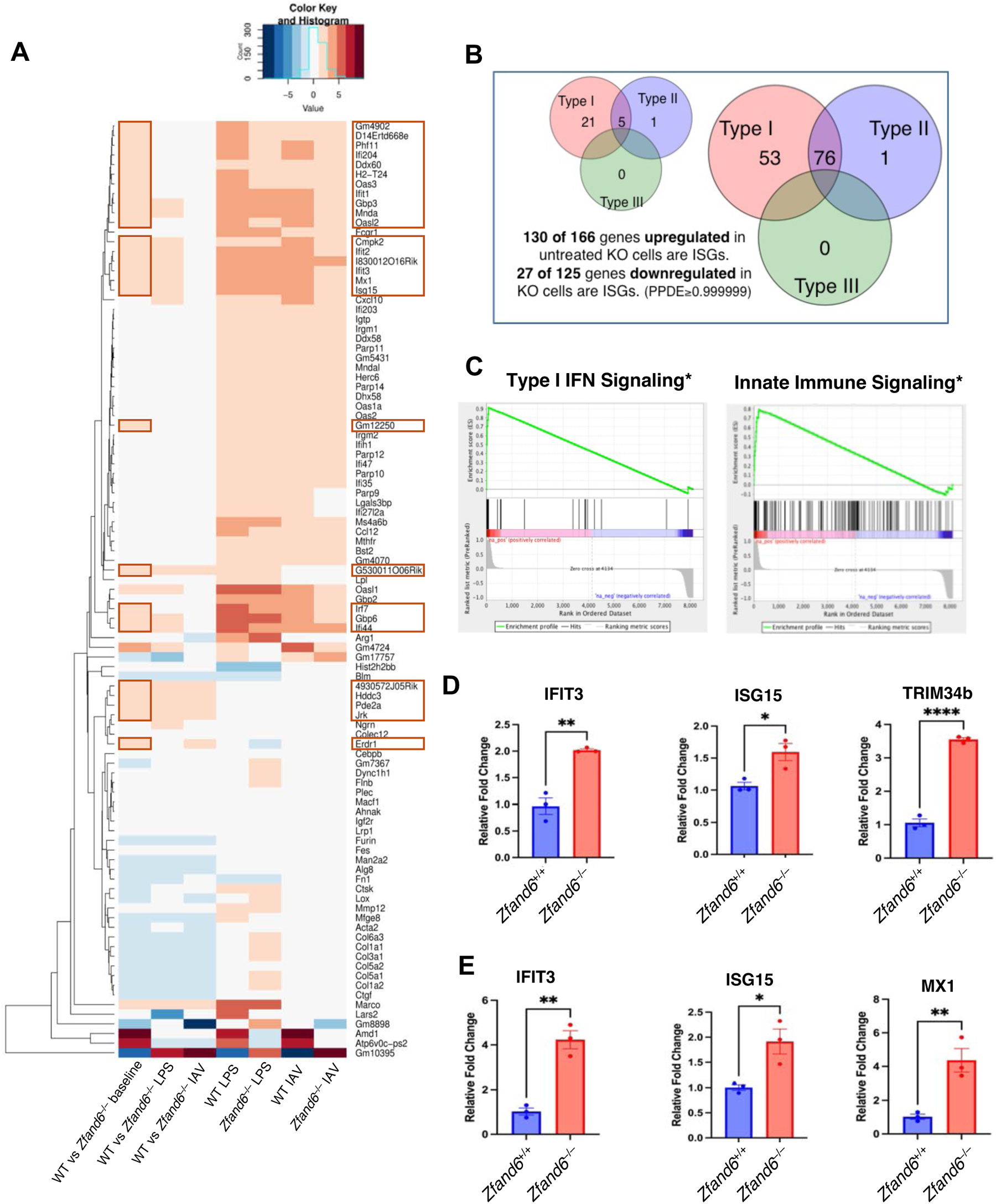
Upregulated ISGs in *Zfand6*^−/−^ BMDMs and MEFs. RNA was isolated from *Zfand6*^+/+^ and *Zfand6*^−/−^ BMDMs for RNA-Seq. **(A)** ISGs were identified from the set of differentially expressed genes (PPDE>0.999999) in KO vs WT cells using Interferome.org. Heat map of selected genes. Genes enclosed in red boxes are ISGs. **(B)** ISGs comprised the majority of genes upregulated in KO cells compared to WT. Venn Diagram of ISGs was generated from the set of genes upregulated in untreated KO vs WT cells using Interferome.org. **(C)** GSEA analysis of RNA-Seq results indicates type I IFN and innate immune signaling pathways are upregulated in *Zfand6*^−/−^ BMDMs. **(D, E)** Upregulation of select ISGs in *Zfand6*^−/−^ BMDMs **(D)** and MEFs **(E)** was confirmed by qRT-PCR (upper panel). Data presented as mean **±**SEM. Unpaired Student’s t-test. *P<0.05, **P<0.05, ****P<0.0001

### *Zfand6*^−/−^ MEFs and BMDMs are resistant to DNA and RNA virus infections

Since ISGs are critical to restrict viral infection and spread^33^, we hypothesized that ZFAND6 deficiency would impart resistance to viral infections. Indeed, *Zfand6*^−/−^ MEFs were broadly resistant to a panel of DNA (mCMV, HSV-1) and RNA (SeV, IAV) viruses (Fig. 2A-F, Extended Data Fig. 2). The resistance to VSV infection was lost when ZFAND6 expression was restored by lentiviral transduction (Extended Data Fig. 2A). Furthermore, infection of *Zfand6*^−/−^ BMDMs and MEFs with Sendai virus (SeV) yielded heightened levels of interferon-β (IFNβ) and interferon-α4 (IFNα4) mRNAs (Fig. 2G-H, Extended Data Fig. 2D) suggesting that increased type I IFN (IFN-I) accounts for the viral resistance in *Zfand6*^−/−^ MEFs and BMDMs.

**Figure 2.**
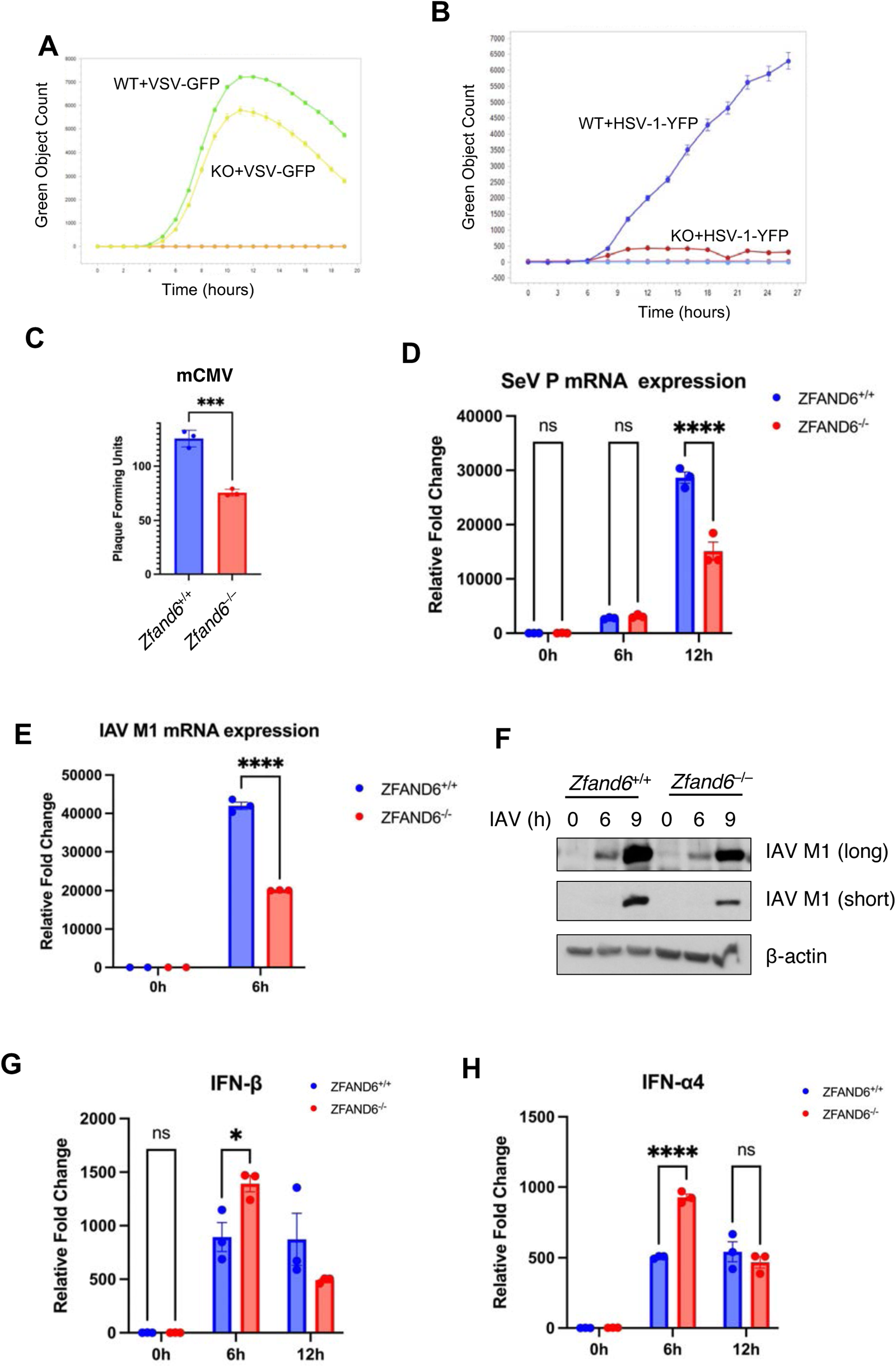
Loss of ZFAND6 imparts resistance to viral infections. **(A)** IncuCyte S3 live-cell imaging of WT and *Zfand6*^−/−^ MEFs infected with VSV-GFP (MOI=1). **(B)** IncuCyte S3 live-cell imaging of WT and *Zfand6*^−/−^ MEFs infected with HSV-1-YFP (MOI=1). (**C)** Plaque assay of WT and *Zfand6*^−/−^ MEFs infected with mCMV (MOI=1). **(D)** qRT-PCR of SeV P in WT and *Zfand6*^−/−^ MEFs infected with SeV (30 HA units). Data presented as mean **±** SEM **(E)** qRT-PCR of IAV M1 in WT and *Zfand6*^−/−^ MEFs infected with IAV (MOI=1). Data presented as mean **±** SEM **(F)** Western blotting of IAV M1 (long exposure on top, short exposure on bottom) and b-actin using lysates from WT and *Zfand6*^−/−^ BMDMs infected with IAV (MOI=1). **(G, H)** qRT-PCR of IFN-β and IFN-a4 in WT and *Zfand6*^−/−^ MEFs infected with SeV (30 HA units) at the indicated time points Data presented as mean **±** SEM. Unpaired Student’s t-test was performed for mCMV infection and 2-way ANOVA with multiple comparisons was performed for SeV and IAV experiments. ***P<0.001, ****P<0.0001, ns=not significant.

### *Zfand6*^−/−^ mice are more susceptible to IAV infection due to impaired myeloid cell activation and diminished IFN signaling

Based on our *in vitro* data, we hypothesized that *Zfand6*^−/−^ mice would be more resistant to an RNA virus infection, therefore we challenged *Zfand6*^−/−^ mice with a low dose of mouse-adapted IAV (A/California/4/2009/H1N1). Surprisingly, mortality was significantly increased in IAV-infected *Zfand6*^−/−^ mice compared to wild-type (WT) mice (Fig. 3A). We did not observe any differences in mortality in *Zfand6*^−/−^ mice challenged with IAV California strain or PR8, thus all subsequent studies were performed with the PR8 strain. *Zfand6*^−/−^ mice infected with IAV PR8 also exhibited more weight loss compared to WT mice (Fig. 3B). We next quantified viral gene expression (*NS1* and *M1*) by real-time qRT-PCR, and found that expression of both *NS1* and *M1* genes was markedly increased in the lungs of *Zfand6*^−/−^ mice at day 7 post-infection (Fig. 3C). The increased viral load instigated more inflammation and tissue damage in the lungs of *Zfand6*^−/−^ mice (Fig. 3D). Therefore, *Zfand6*^−/−^ mice are more susceptible to low doses of IAV.

**Figure 3.**
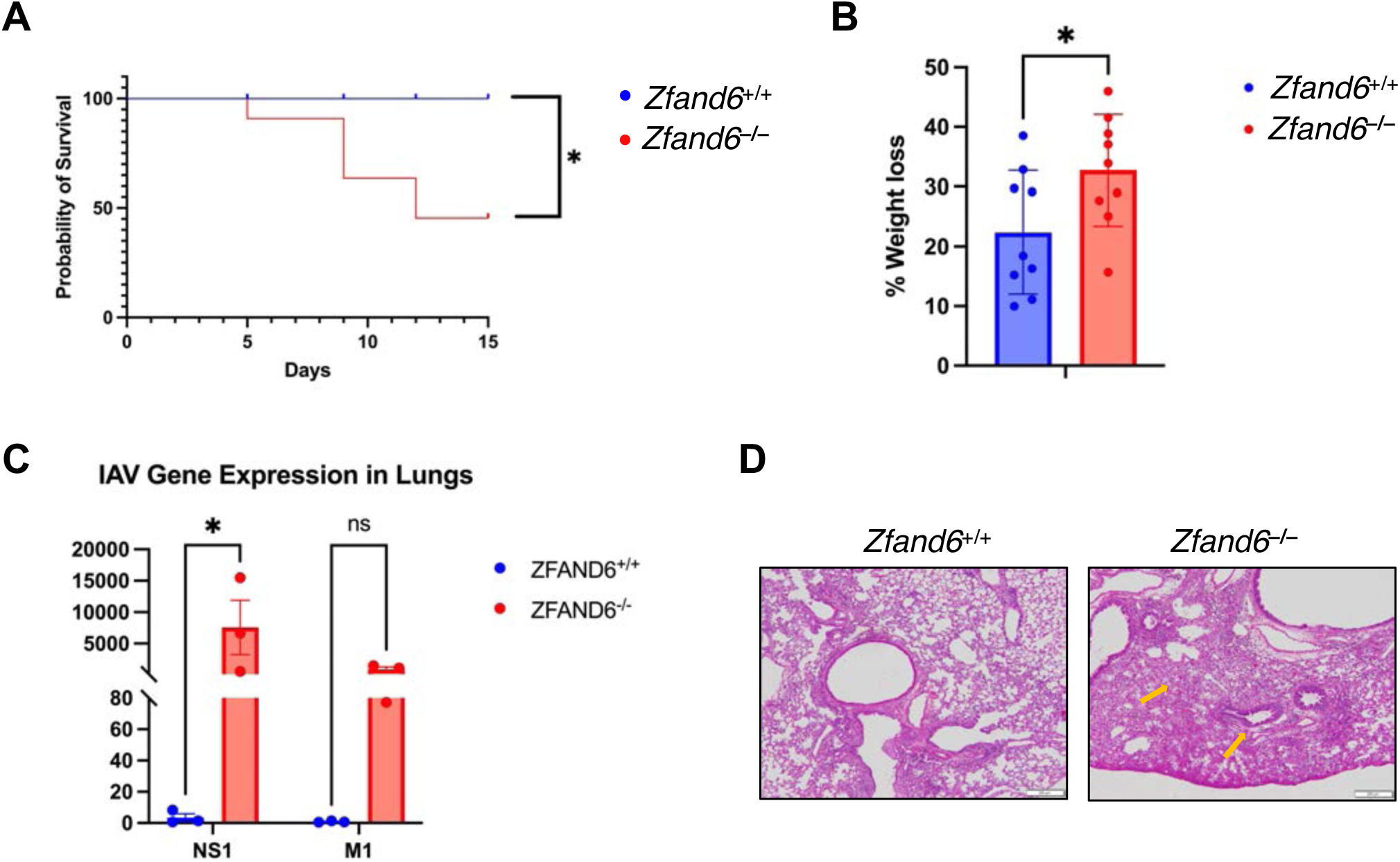
*Zfand6*^−/−^ mice exhibit heightened mortality upon challenge with a sublethal dose of IAV. **(A)** Mice (+/+ n=5, –/– n=11) were infected intranasally with a low dose (0.1 50% mouse lethal dose MLD_50_) of IAV (2009; California; H1N1). Survival was plotted as a Kaplan Meier curve. Wilcoxon-Cox test. *P<0.03 **(B)** Mice (+/+ n=8, -/-n=9) WT and *Zfand6*^−/−^ mice were infected with 1000 focus forming units (FFC) of IAV (PR8) and weight of the mice was monitored daily. The % weight loss was graphed at day 10. Data presented as mean **±** SEM. **(C)** Mice (+/+ n=3, -/-n=3) WT and *Zfand6*^−/−^ mice were infected with 1000 FFC of IAV (PR8) and NS1 and M1 gene expression in lung tissue were detected by qRT-PCR. Data presented as mean **±** SEM. **(D)** H&E stains of lung tissue from WT and *Zfand6*^−/−^ mice at day 10 p.i. with 1000 FFC IAV. Unpaired Student’s t-test. *P<0.05, ns=not significant.

To examine type I IFN responses *in vivo*, *Zfand6*^−/−^ mice were crossed with interferon-stimulated response element (ISRE) reporter mice (Mx1-GFP)^34^. In these mice, GFP expression can be used to track type I IFN responses at the single-cell level. *Zfand6*^−/−^ x Mx1-GFP mice were infected with IAV and flow cytometry was used to examine GFP+ cell populations in the lung. Surprisingly, total GFP+ cells were decreased in the lungs of IAV-infected *Zfand6*^−/−^ x Mx1-GFP mice (Fig. 4A). GFP expression was also significantly decreased in CD11b+ myeloid cells and alveolar macrophages (Fig. 4B-C). Myeloid cell activation was impaired *in vivo* in *Zfand6*^−/−^ mice since CD11b^+^CD80^hi^ and CD11b^+^MHC-II^+^ cells were decreased in IAV-infected lungs (Fig. 4D, Extended Data Fig. 3A-B). Similar results were obtained with alveolar macrophages (Fig. 4E). Finally, GFP expression was also decreased in lung endothelial cells (CD45-EPCAM-) in *Zfand6*^−/−^ mice (Fig. 4F), likely due to impaired type I IFN production by myeloid cells in *Zfand6*^−/−^ mice. Together, these results suggest that the type I IFN response is impaired *in vivo* in *Zfand6*^−/−^ mice thus rendering these mice more sensitive to IAV infection.

**Figure 4.**
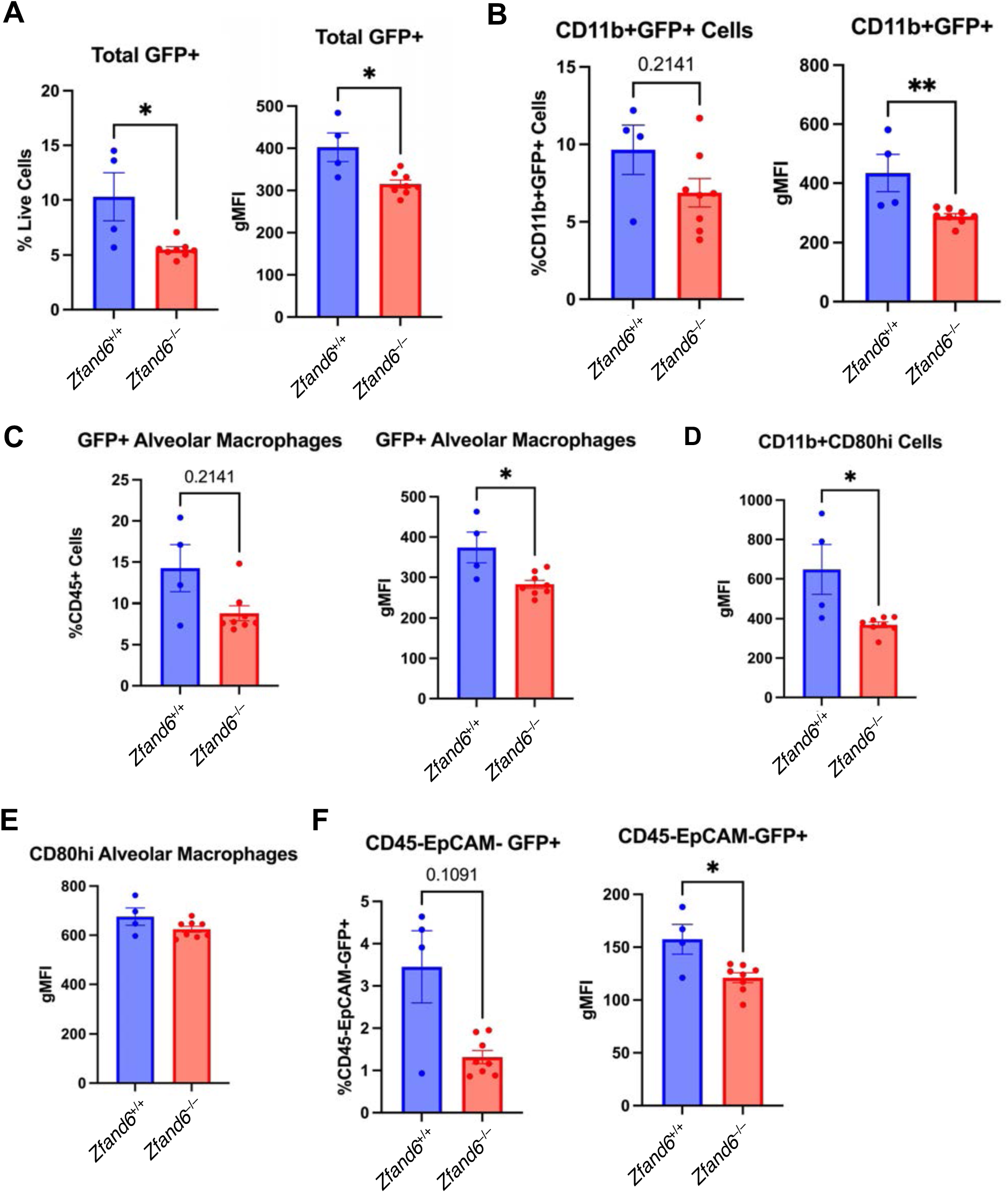
Impaired type I IFN response and myeloid cell differentiation in IAV-infected *Zfand6*^−/−^ mice. **(A-F)** *Zfand6^+/+^MX1-GFP^+/+^* and *Zfand6^-/-^MX1-GFP^+/+^*mice were infected with 1000 FFC of IAV (PR8). Flow cytometry was performed to detect total GFP+ cells in lungs of IAV-infected mice **(A)**, GFP+ cells in CD11b+ myeloid cells **(B)**, alveolar macrophages. Alveolar Macrophages are defined as CD11c^+^SiglecF^+^ cells. **(C)** and CD45-EpCAM-cells (lung endothelial cells) **(F)**. Flow cytometry was also used to detect CD11b+CD80^hi^ cells **(D)** and CD80^hi^ alveolar macrophages **(E)**. Mann-Whitney test. Data presented as mean **±** SEM. *P<0.05, **P<0.05, ns=not significant.

To further examine the mechanisms underlying the impairment in myeloid cell activation, we examined activation markers in WT and *Zfand6*^−/−^ BMDMs. Although no difference was noted in %CD11b^+^F4/80^+^ cells in *Zfand6*^−/−^ BMDMs (data not shown), the % of CD11b^+^F4/80^+^CD86^hi^ macrophages trended lower (Extended Data Fig. 3C). We also generated ZFAND6 knockout THP-1 monocytic cells using CRISPR/Cas9 (Extended Data Fig. 3D). A previous study showed that cIAP1-dependent TRAF2 degradation is required for the differentiation and full activation of macrophages^35^. WT and ZFAND6 KO THP-1 cells were treated with the phorbol ester PMA (phorbol 12-myristate 13-acetate) to induce differentiation and TRAF2 expression was examined by western blotting. Interestingly, TRAF2 degradation was impaired in ZFAND6 KO THP-1 cells treated with PMA (Extended Data Fig. 3E). Therefore, ZFAND6 appears to be essential for TRAF2 degradation during macrophage differentiation, and in the absence of ZFAND6 macrophage activation markers are diminished.

### Loss of ZFAND6 primes the cGAS-STING pathway for activation

The cGAS-STING pathway induces type I IFN expression in response to DNA virus infection^36^, but can also be activated by RNA viruses such as VSV and SARS-CoV2^37, 38^. Therefore, we investigated whether STING contributed to the viral resistance of *Zfand6*^−/−^ cells. The basal expression of ISGs was downregulated in *Zfand6*^−/−^ MEFs by inhibition of STING with the small molecule inhibitor H-151 and rescued the sensitivity to VSV-GFP infection (Fig. 5A, B). Thus, the increased ISG expression and resistance to viral infection in *Zfand6*^−/−^ knockout cells are mediated by STING.

**Figure 5.**
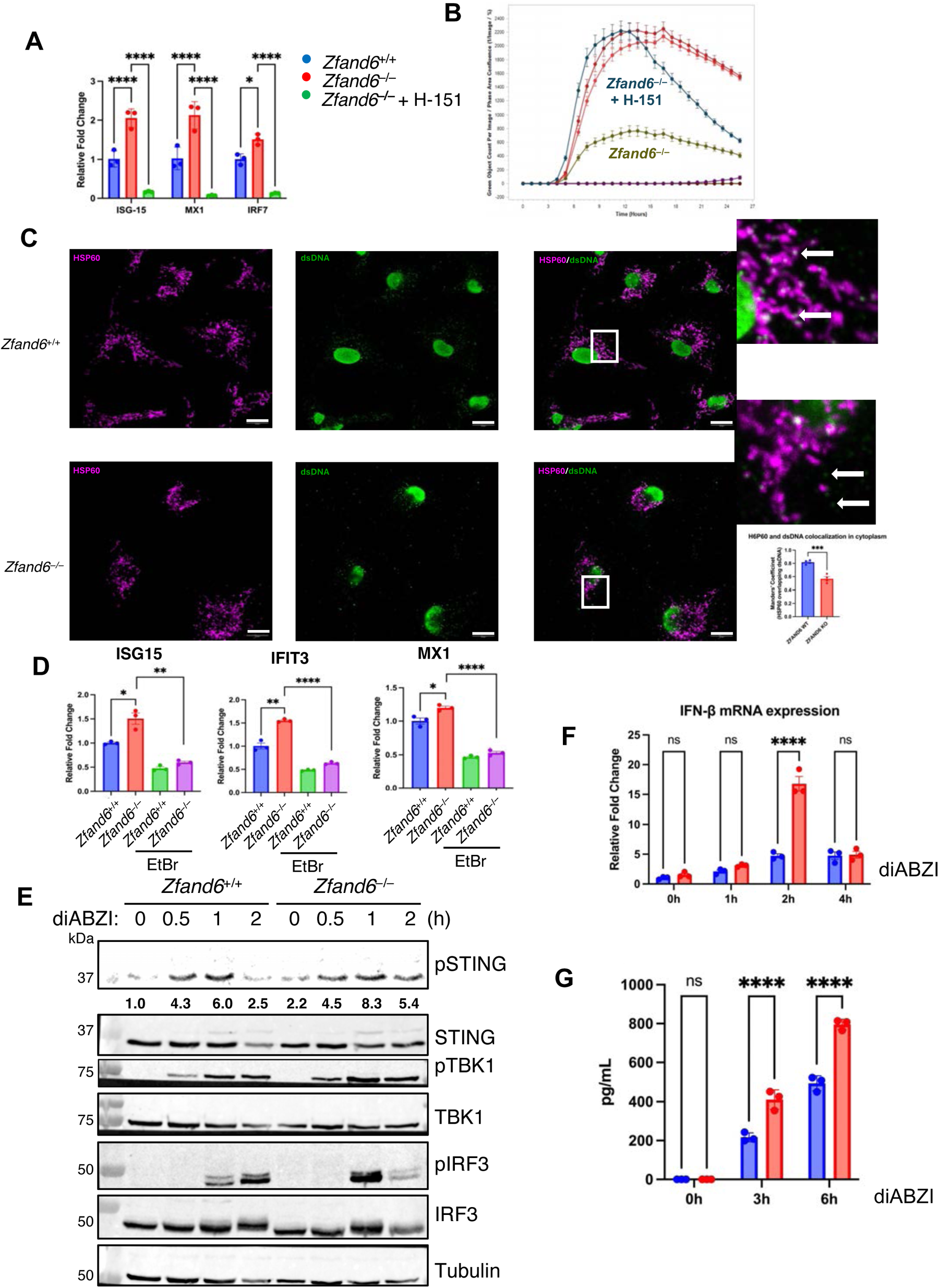
STING activation is sensitized in *Zfand6*^−/−^ BMDMs. **(A)** qRT-PCR of the indicated ISGs in WT and *Zfand6*^−/−^ BMDMs basally and after STING inhibition with 1 μM H-151 for 16 hrs. **(B)** IncuCyte S3 live-cell imaging of WT and Z*fand6*^−/−^ MEFs infected with VSV-GFP and treated with either vehicle or 1 μM H-151. **(C)** Confocal immunofluorescence microscopy with primary antibodies against dsDNA and the mitochondrial marker HSP60 with fluorescently conjugated secondary antibodies in WT and *Zfand6*^−/−^ BMDMs. Quantification of HSP60-dsDNA colocalization reveals decreased dsDNA in the mitochondria in *Zfand6*^−/−^ BMDMs. Scale bars represent 10 μm **(D)** WT and *Zfand6*^−/−^ BMDMs were treated with 150 ng/mL EtBr for six days and qRT-PCR was performed for the indicated ISGs. **(E)** Western blotting was performed using protein lysates from WT and *Zfand6^−/−^*BMDMs stimulated with 1 μM diABZI compound 3 for the indicated times. (**F)** qRT-PCR of IFN-β in WT and *Zfand6*^−/−^ BMDMs upon diABZI stimulation at the indicated time points. **(G)** IFN-β ELISA using supernatants from WT and *Zfand6*^−/−^ BMDMs upon diABZI stimulation at the indicated times. Unpaired Student’s t-test. Data presented as mean **±** SEM. *P<0.05, **P<0.01, ****P<0.0001, ns=not significant.

Since leakage of mitochondrial DNA (mtDNA) activates the cGAS-STING pathway^20, 21^, we next determined whether mtDNA DNA leakage was driving STING activation. To determine whether mtDNA leakage was promoting cGAS-STING activation in *Zfand6^−/−^*BMDMs, we performed confocal microscopy imaging studies by staining for dsDNA and the inner mitochondrial membrane protein HSP60. *Zfand6^−/−^*BMDMs had significantly lower levels of dsDNA in the mitochondria, which were increased in the cytoplasm (Fig. 5C), suggesting mtDNA leakage into the cytoplasm. To examine if leakage of mtDNA into the cytoplasm was driving the activation of the cGAS-STING pathway, we treated cells with low-dose ethidium bromide (EtBr) to deplete mtDNA^21, 39, 40^. Indeed, depletion of mtDNA with low-dose EtBr downregulated the basal expression of ISGs in *Zfand6*^−/−^ BMDMs and MEFs (Fig. 5D and data not shown), suggesting that mtDNA was mediating STING activation of ISGs. To determine if mtDNA leakage could prime the cGAS-STING pathway for activation, we activated STING in WT and *Zfand6*^−/−^ BMDMs using the small molecule agonist diABZI compound 3. *Zfand6*^−/−^ BMDMs exhibited more robust activation of the STING pathway and produced elevated levels of IFNβ as compared to WT BMDMs (Fig. 5E-G).

### Damaged mitochondria accumulate in *Zfand6*^−/−^ MEFs and BMDMs

Since damaged mitochondria have been linked to the leakage of mtDNA^2, 20, 31^, we investigated whether mitochondrial homeostasis was perturbed which in turn caused the mtDNA leakage in *Zfand6*^−/−^ BMDMs. Tetramethyl rhodamine Methyl Ester (TMRM) and MitoTracker Green staining revealed significantly increased numbers of depolarized mitochondria with lower mass in *Zfand6*^−/−^ BMDMs and MEFs (Fig. 6A) indicative of an increased number of fragmented and dysfunctional mitochondria. Furthermore, in agreement with a previous study demonstrating increased ROS levels in ZFAND6 knockout breast cancer cells^15^, *Zfand6*^−/−^ BMDMs similarly had a significant increase in the levels of ROS (Fig. 6B). *Zfand6*^−/−^ BMDMs and MEFs treated with LPS also had significantly more ROS than WT controls that was maintained at high levels throughout the time course of the treatments (Extended Data Fig. 5). Together, these data suggest that mitochondria are dysfunctional in *Zfand6*^−/−^ MEFs and BMDMs.

**Figure 6.**
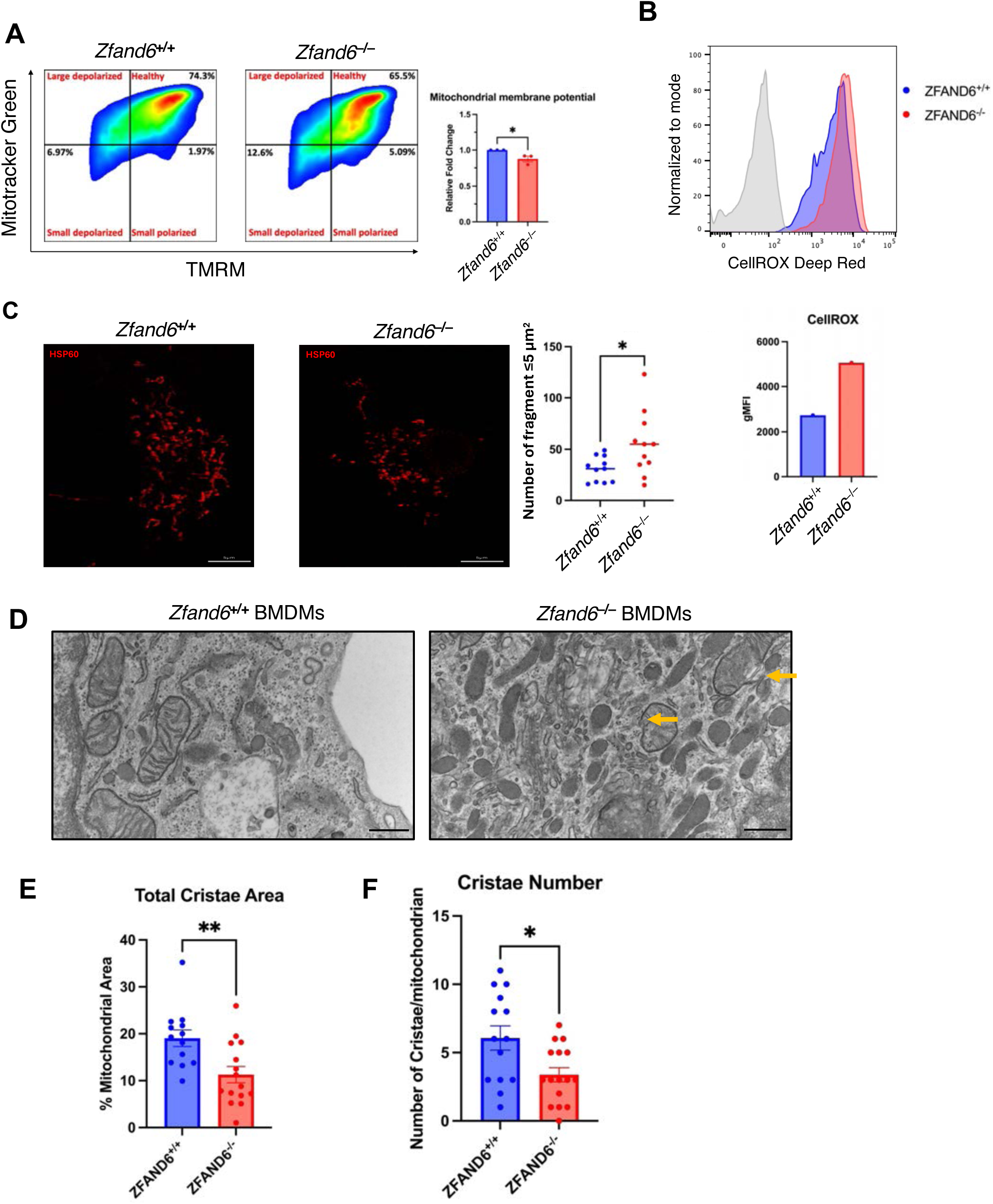
*Zfand6*^−/−^ BMDMs have dysregulated mitochondrial homeostasis. **A**) Flow cytometric analysis of WT and *Zfand6*^−/−^ BMDMs stained with 1:1000 TMRM and 100 nM MitoTracker Green. **B)** Flow cytometric analysis of WT and *Zfand6*^−/−^ BMDMs stained with 5 μM CellROX to measure ROS. CellROX staining was quantified by measuring the geometric mean fluorescence intensity in WT and *Zfand6*^−/−^ BMDMs. **C)** STED microscopy was performed using primary antibody against HSP60 with fluorescently conjugated secondary antibody in WT and *Zfand6*^−/−^ BMDMs. Scale bars represent 5 μm **D)** Transmission electron microscopy analysis of WT and *Zfand6*^−/−^ BMDMs reveals compromised mitochondrial membrane integrity in disrupted cristae networks in *Zfand6*^−/−^ BMDMs Scale bars represent 400 nm. **E, F)** The total mitochondrial cristae area and number were quantified from TEM micrographs using ImageJ. Data presented as mean **±** SEM. Unpaired Student’s t-test. *P<0.05, **P<0.01.

To directly visualize mitochondrial morphology, we imaged mitochondria at super-resolution using stimulated emission depletion (STED) confocal microscopy. Interestingly, STED imaging revealed an increase in fragmented mitochondria in *Zfand6*^−/−^ BMDMs (Fig. 6C). To examine mitochondrial morphology at higher resolution, we next performed transmission electron microscopy (TEM) using WT and *Zfand6*^−/−^ BMDMs. Surprisingly, TEM revealed a significant disruption of the cristae network, loss of mitochondrial membrane integrity and accumulation of electron-dense mitochondria in *Zfand6*^−/−^ BMDMs (Fig. 6D-F), consistent with phenotypic changes associated with impaired mitophagy^31^.

### ZFAND6 is required for mitophagy

To address the mechanism underlying the accumulation of damaged mitochondria in the absence of ZFAND6, we examined if the clearance of mitochondria by mitophagy was impaired. To quantify mitophagy, we employed mtKeima, a fluorescent reporter protein which undergoes a large shift in its excitation peak from 440 nm to 586 nm (pH 7 to pH 4) when mitochondria are delivered to lysosomes^41^. *Zfand6*^−/−^ MEFs stably expressing mtKeima exhibited a defect in mitophagy induction upon treatment with oligomycin A, a potent inhibitor of ATP synthase, and antimycin A, an inhibitor of the electron transport chain (ETC) (O/A) (Fig. 7A). We next performed confocal microscopy with fluorescently conjugated antibodies to the lysosomal marker LAMP1 and the mitochondrial marker HSP60 in WT and *Zfand6*^−/−^ BMDMs treated with O/A. There was an impairment in mitophagosomes (mitochondria in autophagosomes) in *Zfand6*^−/−^ BMDMs with acute mitochondrial damage (Fig. 7B). To further examine if mitophagy was impaired in BMDMs, we treated *Zfand6^+/+^* and *Zfand6*^−/−^ BMDMs with O/A and observed mitophagosomes using TEM. There was a significant decrease in mitophagosomes in *Zfand6*^−/−^ BMDMs upon O/A treatment (Fig. 7C). Furthermore, upon treatment with O/A, *Zfand6*^−/−^ BMDMs exhibited impaired degradation of the mitochondrial protein MTCO2 further supporting a defect in mitophagy (Fig. 7D). Together, these data indicate that ZFAND6 functions as an essential regulator of mitophagy.

**Figure 7.**
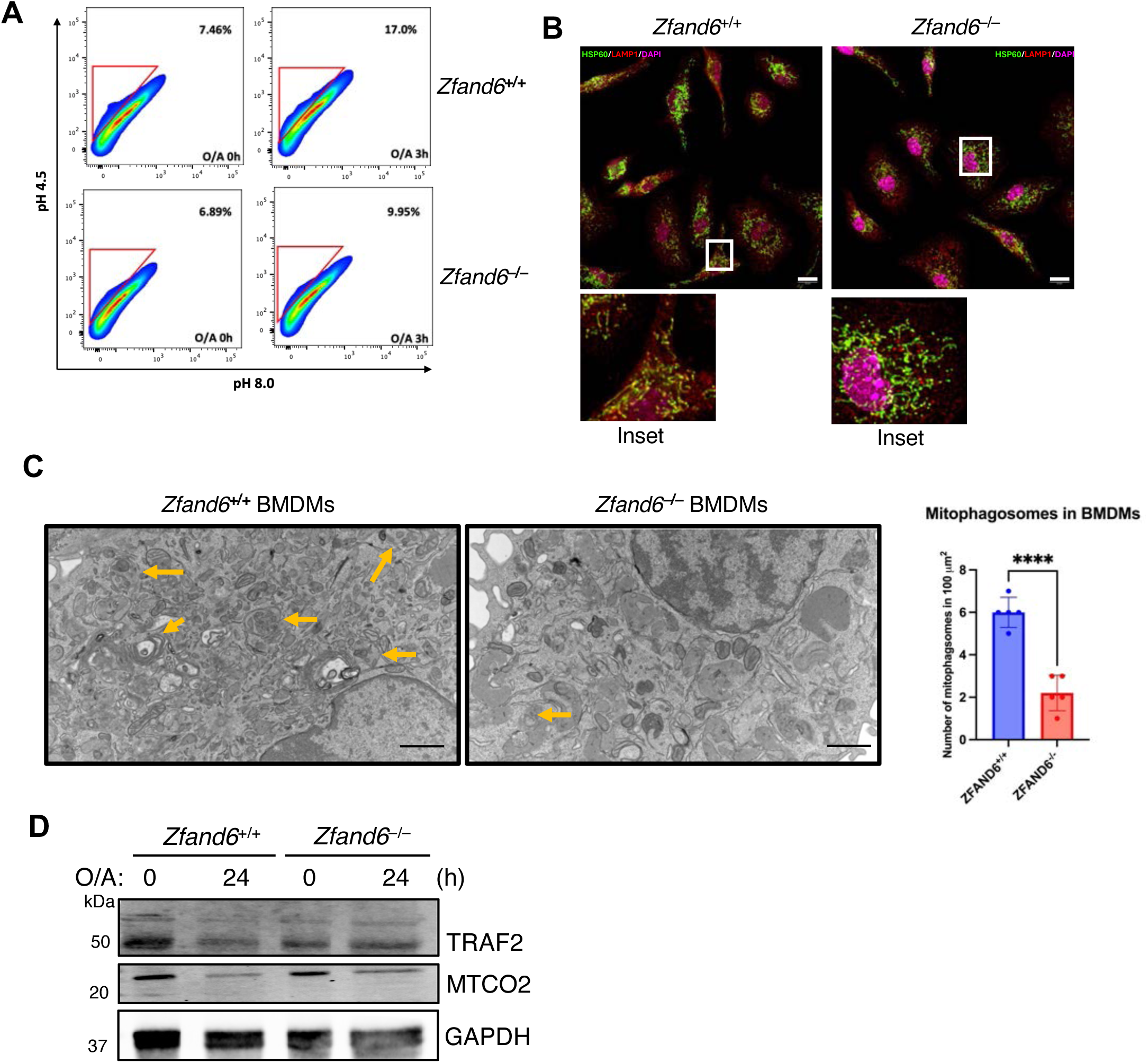
Impaired mitophagy in *Zfand6*^−/−^ BMDMs. **(A)** mtKeima flow cytometry assay using WT and *Zfand6*^−/−^ MEFs either untreated or treated with 5 μM oligomycin A and 1 μM antimycin A for 3 hr. **B)** Confocal microscopy with fluorescently conjugated antibodies to the lysosomal marker LAMP1 and the mitochondrial marker HSP60 in WT and *Zfand6*^−/−^ BMDMs treated with 5 μM oligomycin A and 1 μM antimycin A in presence of 100 μM leupeptin. Scale bars represent 10 μm **C)** Transmission electron microscopy of WT and *Zfand6*^−/−^ BMDMs treated with 5 μM oligomycin A and 1 μM antimycin A in the presence of 20 nM Bafilomycin A1. Scale bars represent 1 μm. Data presented as mean **±** SEM. Unpaired Student’s t-test. ****P<0.0001 **(D)** Western blotting of the indicated proteins from WT and *Zfand6*^−/−^ BMDMs treated with O/A for 24 hr.

### ZFAND6 is required for TRAF2-cIAP1 interaction upon mitophagy induction

The E3 ligase TRAF2 has been implicated in mitophagy and has been shown to interact with ZFAND6^14, 42^. Our earlier experiment also indicated that TRAF2 degradation was impaired in ZFAND6 KO THP-1 cells treated with PMA to induce macrophage differentiation (Extended Data Fig. 3E). Therefore, we hypothesized that ZFAND6 regulated mitophagy through TRAF2. We found that TRAF2 was degraded in response to acute mitochondrial damage caused by O/A treatment, but this was impaired in *Zfand6*^−/−^ BMDMs (Fig. 7D). In the initial steps of mitophagy, damaged mitochondria are conjugated with polyubiquitin chains by E3 ubiquitin ligases such as Parkin^43^ or TRAF2/cIAP^42, 43, 44^. The kinase PINK1 phosphorylates Ub on Ser65 to activate Parkin E3 activity^26, 45^. Since *Zfand6*^−/−^ BMDMs lacked mitophagosomes, it is likely that the early steps of mitophagy were impaired in ZFAND6 knockout cells. Indeed, *Zfand6*^−/−^ BMDMs had significantly decreased levels of p-Ub when mitochondria were damaged in response to O/A treatment as compared to WT BMDMs (Fig. 8A). These data suggest that ZFAND6 promotes early steps of mitophagy upon acute damage of mitochondria.

**Figure 8.**
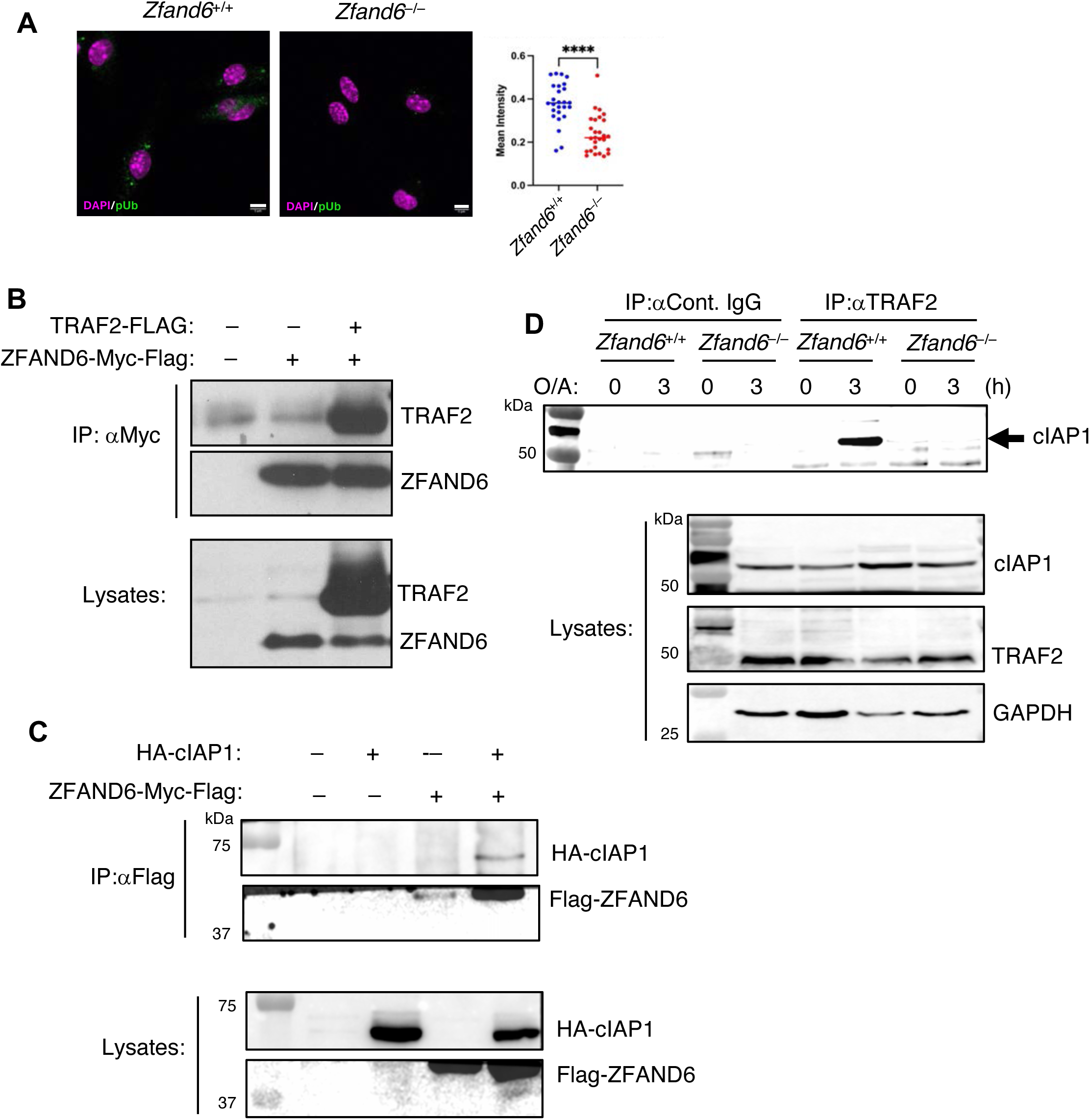
ZFAND6 is required for TRAF2-cIAP1 interactions during acute mitochondrial damage. **A)** Confocal microscopy with fluorescently conjugated antibodies to pUb using WT and *Zfand6*^−/−^ BMDMs treated with 5 μM oligomycin A and 1 μM antimycin A for 3 hr. in the presence of 100 μM leupeptin. Scale bars represent 5 μm. Data presented as mean **±** SEM. (**B, C)** Co-IP assays were performed with lysates from 293T cells transfected with the indicated plasmids. Immunoprecipitations and western blotting were performed with the indicated antibodies to examine ZFAND6-TRAF2 **(B)** and ZFAND6-cIAP1 **(C)** interactions. **(D)** Co-IP assay was performed with lysates from WT and *Zfand6*^−/−^ BMDMs treated with 5 μM oligomycin A and 1 μM antimycin A for the indicated times. IPs were performed with either isotype control antibody or TRAF2 antibody. Western blotting was performed with the indicated antibodies. Unpaired Student’s t-test. ****P<0.0001.

We hypothesized that ZFAND6 was regulating mitophagy through the TRAF2-cIAP complex that was previously shown to induce mitophagy^44^. Thus, we performed co-immunoprecipitation (co-IP) assays to examine if ZFAND6 interacts with TRAF2 and cIAP1. 293T cells were transfected with either empty vector or plasmids encoding tagged versions of ZFAND6, TRAF2 and cIAP1. Our co-IP assay indicated that ZFAND6 could interact with TRAF2 (Fig. 8B), consistent with a previous study^14^. A query of BioGRID for putative ZFAND6 binding proteins indicated that cIAP1 (also known as BIRC2) could potentially be a ZFAND6 binding partner. Two independent mass spectrometry screens aimed at elucidating the human interactome both found potential ZFAND6-cIAP1 interactions^46, 47^. Indeed, our co-IP assay validated the ZFAND6-cIAP1 interaction (Fig. 8C). Given the known interaction between ZFAND6 and TRAF2^14^, the role of TRAF2 as a positive regulator of mitophagy in cardiac cells^31^, and TRAF2 functioning with cIAP1 to regulate mitophagy^42^, we next investigated if ZFAND6 was required for the TRAF2-cIAP1 interaction induced by mitochondrial damage. *Zfand6*^+/+^ and *Zfand6*^−/−^ BMDMs were treated with O/A to induce acute mitochondrial damage and co-IPs were performed to examine TRAF2-cIAP1 interactions. In WT BMDMs, TRAF2 and cIAP1 inducibly interacted in response to O/A treatment; however, the interaction between TRAF2 and cIAP1 was completely impaired in *Zfand6*^−/−^ BMDMs (Fig. 8D). Together, these results suggest that ZFAND6 plays an essential role in supporting TRAF2-cIAP1 interactions during mitophagy and is likely an adaptor/subunit of a TRAF2-cIAP ubiquitin ligase complex.

ZFAND6 contains an A20-like ZF domain at its N-terminus that has sequence similarity with A20 ZF4, and a C-terminal AN1 ZF domain (Extended Data Fig. 6A). We examined if ZFAND6, and either the A20-like ZF or AN1 ZF, could interact with K48, K63 or linear M1-linked polyubiquitin chains. Point mutations were generated for the A20-like ZF domain (C30A) and the AN1 domain (C152A). Interestingly, ZFAND6 could bind K48, K63 and linear polyUb chains, and mutation of the A20-like ZF domain abrogated the Ub binding activity of ZFAND6 (Extended Data Fig. 6B). Therefore, similar to A20, ZFAND6 binds to polyUb chains through its A20-like ZF domain.

## Discussion

Recent studies have underscored the importance of mitochondrial homeostasis in preventing aberrant innate immune signaling and inflammation^2, 21, 31, 48, 49, 50^. As such, inflammatory DAMPs leaked from damaged mitochondria have been linked to several autoimmune disorders^51, 52, 53, 54^. Our findings implicate ZFAND6 as a novel regulator of mitophagy through the regulation of the TRAF2-cIAP E3 ligase complex. ZFAND6 has an A20-like ZF and an AN1 ZF domain, and we found that the A20-like ZF domain of ZFAND6 binds to polyUb chains (Extended Data Fig. 6). It will be important to determine the requirements of the ZFAND6 A20-like ZF and AN1 ZF domains in the induction of mitophagy. We speculate that ZFAND6 will require its Ub binding domain (A20-like ZF) to promote mitophagy. ZFAND6 may potentially bind to ubiquitinated TRAF2 and/or cIAP1 when activated and function as an adaptor, or alternatively ZFAND6 may bind to ubiquitinated proteins at the outer mitochondrial membrane triggered by membrane depolarization and mitochondrial damage and recruit the TRAF2/cIAP complex. Future studies should investigate these possibilities.

In contrast to *Tnfaip3*^−/−^ (A20 knockout) mice, *Zfand6*^−/−^ mice do not exhibit severe multi-organ inflammation and have normal lifespans (data not shown). Our data reveal fundamental differences in the mechanisms by which A20 and ZFAND6 restrict inflammation. Whereas A20 broadly and directly inhibits inflammatory signaling pathways in negative feedback loops by NF-κB-mediated induction of A20 expression^55^, ZFAND6 has evolved a more specialized function to prevent inflammation triggered by mitochondrial damage/impaired mitochondrial homeostasis. Unlike A20, ZFAND6 expression is not strongly induced by inflammatory stimuli (data not shown) indicating yet another difference in their regulation. Furthermore, ZFAND6 protein is highly expressed in monocytes, peripheral blood mononuclear cells (PBMCs), fetal heart, Islet of Langerhans, placenta, fetal ovary and fetal testis (GeneCards database).

Loss of ZFAND6 triggers elevated levels of ROS, and our work has defined the mechanistic basis for this elevated ROS. *Zfand6*^−/−^ BMDMs exhibited pronounced defects in mitochondrial morphology due to impaired mitophagy, which results in the leakage of mtDNA that subsequently activates the cGAS-STING pathway. While inhibition of STING with a small molecule inhibitor diminished the expression of basal ISGs in *Zfand6*^−/−^ MEFs, we cannot rule out activation of endosomal nucleic acid sensing pathways (i.e., TLR7/9) by mtDNA leakage in macrophages as previously reported^2, 31^. Our results suggest that priming of the cGAS-STING pathway rendered *Zfand6*^−/−^ BMDMs and MEFs resistant to VSV and HSV-1 infections, but not SeV (data not shown). This could be explained by either the potential priming of endosomal TLRs, or the accumulation of mitochondrial antiviral signaling (MAVS) aggregates on damaged mitochondria. Upon activation of RIG-I-like receptors, MAVS aggregates accumulate on the OMM to induce downstream IFN-I signaling^56^. MAVS subsequently induces mitophagy to clear the aggregates, thereby providing a mechanism of negative regulation of IFN-I signaling initiated by RNA virus infections^57, 58^. Impairment of mitophagy in *Zfand6*^−/−^ cells could therefore disrupt the homeostasis of other innate immune pathways. Thus, leakage of mitochondrial DAMPs could prime a broad range of PRRs, leading to more robust activation by pathogens. This provides a likely explanation for the resistance to a wide range of viruses in *Zfand6*^−/−^ cells.

In addition to TLRs, leakage of mtDNA has also been proposed to activate NLRP3 and AIM2 inflammasomes^50, 59^. With the accumulation of damaged mitochondria in *Zfand6*^−/−^ MEFs and BMDMs, it will be interesting to determine if *Zfand6*^−/−^ BMDMs are primed for inflammasome activation. Inflammasomes are implicated in a broad range of diseases including type 2 diabetes (T2D)^60, 61^, and single nucleotide polymorphisms (SNPs) in ZFAND6 have been linked to T2D susceptibility^62, 63^. Future studies should investigate if loss of ZFAND6 in mice leads to reduced glucose tolerance and the development of T2D due to the aberrant activation of the NLRP3 inflammasome.

Our data indicate that ZFAND6 is required for cIAP1-mediated TRAF2 degradation which is necessary for full activation of macrophages upon differentiation. We also found that ZFAND6 is essential for TRAF2 degradation triggered by ER stress inducers (data not shown). Furthermore, TRAF2-cIAP1 interactions triggered by acute mitochondrial damage are completely dependent on ZFAND6. Therefore, ZFAND6 likely bridges the TRAF2-cIAP1 interaction as an adaptor protein/subunit of a TRAF2-cIAP E3 ubiquitin ligase complex. The TRAF2-TRAF3-cIAP E3 ligase complex is a key regulator of NF-1B-inducing kinase (NIK) degradation and non-canonical NF-κB signaling, cell survival, TLR signaling and lymphoid cell maturation^29, 64, 65, 66, 67, 68^. Our data suggest that ZFAND6 is a critical regulator of the TRAF2-cIAP1/2 complex in the context of mitophagy as well as macrophage activation. It remains an open question whether ZFAND6 regulates the TRAF2-cIAP complex in non-canonical NF-κB signaling and/or TNF-induced NF-κB signaling and cell survival. Future studies should investigate how loss of ZFAND6 and the subsequent disruption of the cIAP-TRAF2 complex impacts the broader immune response to pathogens *in vivo*.

## Methods

### Generation of *Zfand6*^−/−^ mice

*Zfand6*^−/−^ mice were generated by Ingenious Targeting Laboratory (Stony Brook, NY). A targeting vector was created using a neomycin (Neo) resistance cassette to replace *Zfand6* exons 2-4. The construct was designed such that the short homology arm extends 2.5 kb to the Neo cassette and the long homology arm extends 5.5 kb away. The targeting vector was electroporated into C57BL/6Nx129/SvEv embryonic stem cells and correctly targeted clones were identified by southern blotting. ES clones were microinjected into C57BL/6N blastocysts to generate chimeric mice. Germline transmission of the disrupted allele was obtained by crossing male chimeras with WT C57BL/6N female mice. Homozygous knockout animals were born at Mendelian ratios after crosses between heterozygous mice. The Neo cassette, flanked by LoxP sites, was deleted in the germ line of *Zfand6*^−/−^ mice by crossing with Ella-Cre mice (strain# 003724; Jackson Laboratory). *Zfand6*^−/−^ mice were backcrossed onto a C57BL/6N genetic background for 10 generations. *Zfand6*^−/−^ mice were also crossed with Mx1-GFP mice (kind gift from Adolfo Garcia-Sastre; Icahn School of Medicine at Mount Sinai).

### Cell culture, plasmids and reagents

Bone marrow-derived macrophages (BMDMs) were generated as described and cultured in DMEM media^69^. Murine embryonic fibroblasts (MEFs) were generated as described and immortalized by transfection of SV40 Large T antigen^70^. MEFs were cultured in DMEM media supplemented with 10% fetal bovine serum (FBS) (Cytiva Cat#SH30071.03) and 1% of 10,000U/mL Penicillin Streptomycin (P/S) (Gibco Cat#15140122). THP-1 cells were purchased from ATCC and cultured in RPMI media supplemented with 10% FBS and 1% P/S. 293T cells were cultured in DMEM media supplemented with 10% FBS and 1% P/S and transfected with plasmids using GenJet (SignaGen; SL100489). pHAGE-mt-mKeima was a gift from Richard Youle (Addgene plasmid# 131626; http://n2t.net/addgene:131626 ; RRID:Addgene_131626). ZFAND6-Flag-Myc was purchased from Origene. pEBB-HA cIAP1 was a gift from Colin Duckett (Addgene plasmid# 38232; http://n2t.net/addgene:38232; RRID:Addgene_38232). lentiCRISPRv2 was a gift from Feng Zhang (Addgene plasmid# 52691 ; http://n2t.net/addgene:52691; RRID:Addgene_52691). pUltra-Hot was a gift from Malcolm Moore (Addgene plasmid# 24130 ; http://n2t.net/addgene:24130 ; RRID:Addgene_24130). pUltra-Hot-ZFAND6 was generated by subcloning a ZFAND6 PCR product digested with Xba1 (New England Biolabs; R0145S) and BamH1-HF (New England Biolabs; R3136S) into pUltra-Hot. Flag-TRAF2 was a gift from Ze’ev Ronai (Sanford Burnham Prebys). psPAX2 was a gift from Didier Trono (Addgene plasmid# 12260 ; http://n2t.net/addgene:12260; RRID:Addgene_12260). Oligomycin A (cat.#: 75351), antimycin A (cat.#: A8674) and PMA (cat.#: p1585) were purchased from Millipore-Sigma. diABZI compound 3 (cat.#: tlrl-diabzi), LPS (cat.#: tlrl-3pelps) and H-151 (cat.# inh-h151) were purchased from InvivoGen. Recombinant K48, K63 and linear ubiquitin chains were purchased from Boston Biochem.

### Lentivirus and stable cell line generation

2.5×10^6^ Lenti-X 293T cells (Takara; 632180) were seeded into a 10-cm dish and transfected with 6:3:1 ratios of lentiviral plasmid encoding the gene of interest, psPAX2 and VSV-G using GenJet (SignaGen). The media was changed 6 hrs. after transfection and replaced with 10 mL DMEM supplemented with 10% FBS. Cell supernatant was harvested 48 hrs. post-transfection and filtered with a 0.4 μm PES non-pyrogenic filter. Harvested supernatant was concentrated 100x using the Lenti-X Concentrator (Takara; 631231) and resuspended in complete DMEM and stored at -80°C. For stable cell line generation, cells were seeded in a 6-well plate and transduced with 10 μL of concentrated lentivirus in the presence of 10 μg/mL polybrene (Millipore-Sigma; H9268) and replaced with fresh media 24 hrs. post-transduction. Cells were then either selected using antibiotic selection or sorted using flow cytometry if transduced with a plasmid encoding a fluorescent protein 48 hrs. post-transduction.

### Virus infections

VSV-GFP was obtained from Dr. Glen Barber (University of Miami). SeV (Cantell strain) was purchased from Charles River Laboratories. HSV-1-YFP was a gift from Dr. John Wills (Penn State College of Medicine). For virus infections, cells were serum starved for 1 hr. and then adsorbed with the indicated virus and MOIs in serum-free media for 1 hr. with periodic rocking. Cells were then incubated with complete media and viral loads or gene expression were detected using the indicated methods.

### ZFAND6 CRISPR knockout THP-1 cells

CRISPR/Cas9-mediated knockout of ZFAND6 was performed as described previously^71^. Three guide RNAs (gRNAs) specific for human ZFAND6 were designed using the E-CRISP web server and cloned into lentiCRISPRv2. THP-1 cells were transduced with lentiviruses expressing Cas9 and ZFAND6 gRNAs to generate ZFAND6 knockouts. Transduced cells were selected with puromycin, and then subjected to limiting dilution. Individual clones were screened by western blotting with anti-ZFAND6 (Origene; TA335508).

### *In vivo* IAV infections and lung homogenization

8-15 weeks of age and sex-matched WT and *Zfand6*^−/−^ mice were intranasally infected with either 1/10 of 10^3^ TCID_50_ of IAV H1N1 (A/California/4/2009) or 1000 focus forming units (FFC) of IAV H1N1 (PR8) for the indicated times^72^. Lungs were harvested and subjected to enzymatic digestion using 50 μg/mL Liberase TM (Millipore-Sigma; LBTM-RO) in serum free RPMI at 37°C for 30 min. and then passed through an 18G syringe 12 times and then incubated for an additional 15 min. at 37°C. The enzymatic reaction was terminated by adding 8 mL stop solution (PBS with 2% FBS, 5mM EDTA, pH 8.0). Cells were passed through a 70 μm cell strainer and spun at 500xg and subjected to RBC lysis using RBC Lysis Buffer (BioLegend; 420301) to yield a single-cell suspension. These cells were then further processed for flow cytometric analysis.

### RNA Sequencing

Total RNA was prepared from *Zfand6*^+/+^ and *Zfand6*^−/−^ BMDMs, either untreated, treated with LPS or infected with IAV (PR8) for 24 hr. RNA-Seq and bioinformatics was performed by the Johns Hopkins Sidney Kimmel Comprehensive Cancer Center next-generation sequencing core as previously described^73^. Illumina next-gen sequencing reads were mapped with RSEM, aligned to a mouse genome reference library and analyzed with RSEM, EBSeq, Tophat and Cufflinks. The differential expression analysis was performed with the DEseq R package and GO enrichment with the topGO R package. Gene set enrichment analysis (GSEA) was performed using GSEA (Broad Institute, broadinstitute.org/GSEA). The RNA-Seq data from mouse *Zfand6*^+/+^ and *Zfand6*^−/−^ BMDMs are publicly available in the Sequence Read Archive with an accession code of PRJNA1082378.

### Quantitative Real Time PCR (qRT-PCR)

1×10^6^ BMDMs or 1.5×10^5^ MEFs were seeded in 6-well plates 1 day before the experiment. After the treatments, cells were gently scrapped from the wells and pelleted either to store at -80C° for later analysis or processed immediately. Total RNA was extracted using the RNA Spin II Kit (Macherey Nagel; 740955) using the manufacturer’s instructions, and cDNA was synthesized using M-MLV Reverse Transcriptase (Life Technologies; 28025-013) and Oligo (dT) (Life Technologies; 18418-012). cDNA was then subjected to real-time qPCR using a QuantStudio 3 (ThermoFisher Scientific) to quantify mRNA copies using PowerUP SYBR Green Master Mix (Applied Biosystems; A25742). The mRNA transcripts were quantified as described^74^. Primer sequences for qRT-PCR experiments are listed in Supplementary Table 1.

### Western blotting and Co-immunoprecipitations

Cells were gently scraped in the presence of ice-cold PBS and pelleted. Cells were lysed by resuspending 50 μL/1×10^6^ cells in RIPA lysis buffer. Resuspended cells were rotated for 30 min. at 4°C followed by centrifugation at 16,000x*g* for 15 min. at 4°C and clarified supernatants were collected for SDS-PAGE electrophoresis. Protein concentration was estimated using Pierce^TM^ BCA protein assay kit (ThermoFisher Scientific; 23227), reduced with NuPage sample reducing agent (ThermoFisher Scientific; NP0009) and loaded with NuPage sample loading dye (ThermoFisher Scientific; NP0007) on polyacrylamide gels for electrophoresis. Gels with resolved proteins were transferred to a nitrocellulose membrane for blotting using a Bio-Rad tank transfer system for 90 min. at 110V or a Bio-Rad Turbo Transfer system for 15 min. at manufacturer-recommended voltages. Transferred membranes were blocked with 5% fat-free milk for 1 hr. for total proteins and in SuperBlock^TM^ Blocking Buffer (ThermoFisher Scientific; 37535) for 30 min. for phosphorylated proteins and incubated with primary antibodies in 3% BSA in PBS-T overnight at 4℃ or 1 hr. at room temperature. After incubation with primary antibodies, membranes were washed with PBS-T and incubated with species specific HRP-conjugated secondary antibodies and imaged using an Azure 600 imaging system. Antibodies used for western blotting are: STING Rabbit mAb (Cell Signaling Cat# 13647S), Phospho-STING rabbit anti-mouse mAb (Cell Signaling Cat# 72971S), IRF3 rabbit mAb (Cell Signaling Cat# 4302S), p-IRF3 rabbit mAb (Cell Signaling Cat# 4947S), TBK1 rabbit mAb (Cell Signaling Cat# 3504S), p-TBK1 rabbit mAb (Cell Signaling Cat# 5483S), TRAF2 rabbit pAb (Proteintech Cat# 26846-1-AP), cIAP1 Rabbit pAb (Proteintech Cat# 10022-1-AP), HA rabbit pAb (Proteintech Cat# 51064-2-AP), MTCO2 Rabbit pAb (Proteintech Cat# 55070-1-AP), GAPDH mouse mAb (Proteintech Cat# 60004-1-IG), Flag mouse mAb (Proteintech Cat# 66008-4-Ig), Myc Mouse mAb (Proteintech Cat# 60003-2-Ig) and ZFAND6 rabbit pAb (Origene Cat# TA335508).

*Co-immunoprecipitation:* 500 μg of whole-cell lysate was used for endogenous immunoprecipitations and 300 μg of whole cell lysate was used for immunoprecipitation of transiently expressed proteins. IPs were performed using the Dynabeads Protein G IP kit (ThermoFisher Scientific; 10007D) according to the manufacturer’s instructions.

### Confocal microscopy and super resolution microscopy

7×10^4^ BMDMs were seeded in 8-well chamber slides (Corning; 354118) 24 hrs. prior to the experiment. Cells were fixed with 4% PFA in PBS for 15 min. at room temperature followed by 3x 5-min. washes with PBS. Cells were then permeabilized with 0.5% Triton-X for 5 min. and blocked with 3% BSA in PBS for at least 1 hr. at room temperature. Cells were incubated with appropriate antibodies diluted in 3% BSA in PBS either overnight at 4°C or for 1 hr. at room temperature. After 3x 5-min. washes with PBS, cells were incubated with fluorescently conjugated secondary antibodies diluted in 3% BSA in PBS for 1 hr. at room temperature. Secondary antibodies were washed with 3x 5-min. antibodies and mounted with ProLong Diamond Antifade mountant with DAPI (ThermoFisher Scientific; P36966). Confocal immunofluorescence images were acquired using a Leica SP8 confocal microscope equipped with a 63x 1.4 N.A. objective. Images were deconvolved with Huygens Professional and analyzed using FIJI Image analysis software (NCBI). Antibodies used for confocal microscopy are: Alexa Fluor 647 mouse anti-mouse LAMP1 (Santa Cruz Cat# sc-20011 AF647), dsDNA mouse mAb (Santa Cruz Cat# sc-58749), HSP60 rabbit mAb (Cell Signaling Cat# 12165) and Phospho-Ubiquitin rabbit mAb (Cell Signaling Cat #70973).

### Transmission electron microscopy

3×10^6^ BMDMs were seeded in 6 cm dishes. Samples were fixed by 2.5% glutaraldehyde and 2% paraformaldehyde in 0.1 M phosphate buffer (pH 7.4) and further fixed in 1% osmium tetroxide in 0.1 M phosphate buffer (pH 7.4) for 1 hr. Samples were dehydrated in a graduated ethanol series, acetone, and embedded in LX-112 (Ladd Research, Williston, VT). Thin sections (65nm) were stained with uranyl acetate and lead citrate and viewed with a JEOL JEM1400 Transmission Electron Microscope (JEOL USA Inc., Peabody, MA, USA) in the Penn State College of Medicine TEM Facility (RRID Number: SCR_021200).

### Live-cell imaging

Live-cell imaging was performed using an IncuCyte S3 imaging system (Sartorius) to monitor viral infection as previously reported^75^. Briefly, 3×10^4^ MEFs were seeded in 24-well plates 1 day prior to viral infection. Infected cells were scanned with a 10x objective and viral infection was quantified by counting green objects normalized to phase area.

### Flow cytometry assays

1×10^6^ cells were incubated with 1:1000 Image-iT TMRM Reagent (ThermoFisher Scientific; I34361), 5 μM CellROX Deep Red (ThermoFisher Scientific; C10491), or 100 nM MitoTracker Green FM (ThermoFisher Scientific; M7514) according to the manufacturer’s instructions. Cellular ROS was also detected using the DCFDA/H2DCFDA cellular ROS assay kit according to the manufacturer’s instructions (Abcam; ab113851). To analyze cell populations *ex vivo*, 1×10^6^ purified cells were blocked with TruStain FcX^TM^ (BioLegend; 101320) for 10 min. on ice and then incubated with primary antibodies. To prevent aggregation of polymer dyes, 50 μL/well of BD Horizon^TM^ Brilliant Stain buffer (BD Biosciences; 563794) was included. Cells were washed for 5 min. at 500x*g* and incubated with Live/Dead Fixable viability dye (ThermoFisher Scientific; L34973) and then fixed with IC Fixation buffer (eBiosciences; 00822249) for 20 min. and analyzed using a 17-color BD FACSSymphony flow cytometer. Antibodies used for flow cytometry are: BV650 rat anti-mouse CD45 (Biolegend Cat# 103151), BUV395 Rat anti-mouse CD11b (BD Horizon Cat# 563553), Alexa Fluor 700 Armenian Hamster anti-mouse CD11c (BioLegend Cat#117320), BV 711 Armenian Hamster anti-mouse CD80 (BioLegend Cat# 104743), APC rat anti-mouse Ly-6C (BioLegend Cat# 128015), PE rat anti-mouse I-A/I-E (MHC-II) (BioLegend Cat# 107608), BUV 737 rat anti-mouse Siglec F (Invitrogen Cat# 367-1702-82), Pe-Cy5 rat anti-mouse CD326(Ep-CAM) (BioLegend Cat# 118248) and PE/Fire 810 mouse anti-mouse CD161 (NK1.1) (BioLegend Cat# 156535) Gating strategies for flow cytometry are described in Extended Data Fig. 4.

### Statistical analysis

GraphPad Prism 10 was used for all statistical analysis.

## Acknowledgements

We are grateful to Drs. Sid Balachandran and Glen Barber for VSV-GFP and Dr. John Wills for HSV-1-YFP. We thank Dr. Wasif Khan (University of Miami) for valuable discussions. We acknowledge the Johns Hopkins Sidney Kimmel Comprehensive next-generation sequencing core for RNA-Seq analysis, the Penn State College of Medicine Advanced Light Microscopy Core and the Flow Cytometry Core. The Advanced Light Microscopy Core (RRID: SCR_022526) and Flow Cytometry Core (RRID:SCR_021134), services and instruments used in this project were funded, in part, by the Pennsylvania State University College of Medicine via the Office of the Vice Dean of Research and Graduate Students and the Pennsylvania Department of Health using Tobacco Settlement Funds (CURE). The content is solely the responsibility of the authors and does not necessarily represent the official views of the University or College of Medicine. The Pennsylvania Department of Health specifically disclaims responsibility for any analyses, interpretations, or conclusions. This work was supported, in part, by NIH grants R21 AI137550 and R01 AI162815 (to EWH), the Judy S. Finkelstein Memorial Research Award (to KS), and CRIP (Center for Research on Influenza Pathogenesis), a NIAID-funded Center of Excellence for Influenza Research and Response (CEIRR, contract #75N93021C00014) to A.G.-S.

## Data Availability

All data are available in the main text or supplementary materials. The RNA-Seq data from mouse *Zfand6*^+/+^ and *Zfand6*^−/−^ BMDMs are publicly available in the Sequence Read Archive with an accession code of PRJNA1082378.

## Ethics Declarations

The A.G.-S. laboratory has received research support from GSK, Pfizer, Senhwa Biosciences, Kenall Manufacturing, Blade Therapeutics, Avimex, Johnson & Johnson, Dynavax, 7Hills Pharma, Pharmamar, ImmunityBio, Accurius, Nanocomposix, Hexamer, N-fold LLC, Model Medicines, Atea Pharma, Applied Biological Laboratories, Merck and Castlevax, outside of the reported work. A.G.-S. has consulting agreements for the following companies involving cash and/or stock: Castlevax, Amovir, Vivaldi Biosciences, Contrafect, 7Hills Pharma, Avimex, Pagoda, Accurius, Esperovax, Applied Biological Laboratories, Pharmamar, CureLab Oncology, CureLab Veterinary, Synairgen, Paratus, Pfizer and Prosetta, outside of the reported work. A.G.-S. has been an invited speaker in meeting events organized by Seqirus, Janssen, Abbott, Astrazeneca and Novavax. A.G.-S. is inventor on patents and patent applications on the use of antivirals and vaccines for the treatment and prevention of virus infections and cancer, owned by the Icahn School of Medicine at Mount Sinai, New York, outside of the reported work. A.P. has consulted for GlaxoSmithKline. The other authors declare no competing interests.

