## Extended Figures for "ZFAND6 is a subunit of a TRAF2-cIAP E3 ubiquitin ligase complex essential for mitophagy"

**Extended Data Fig. 1**

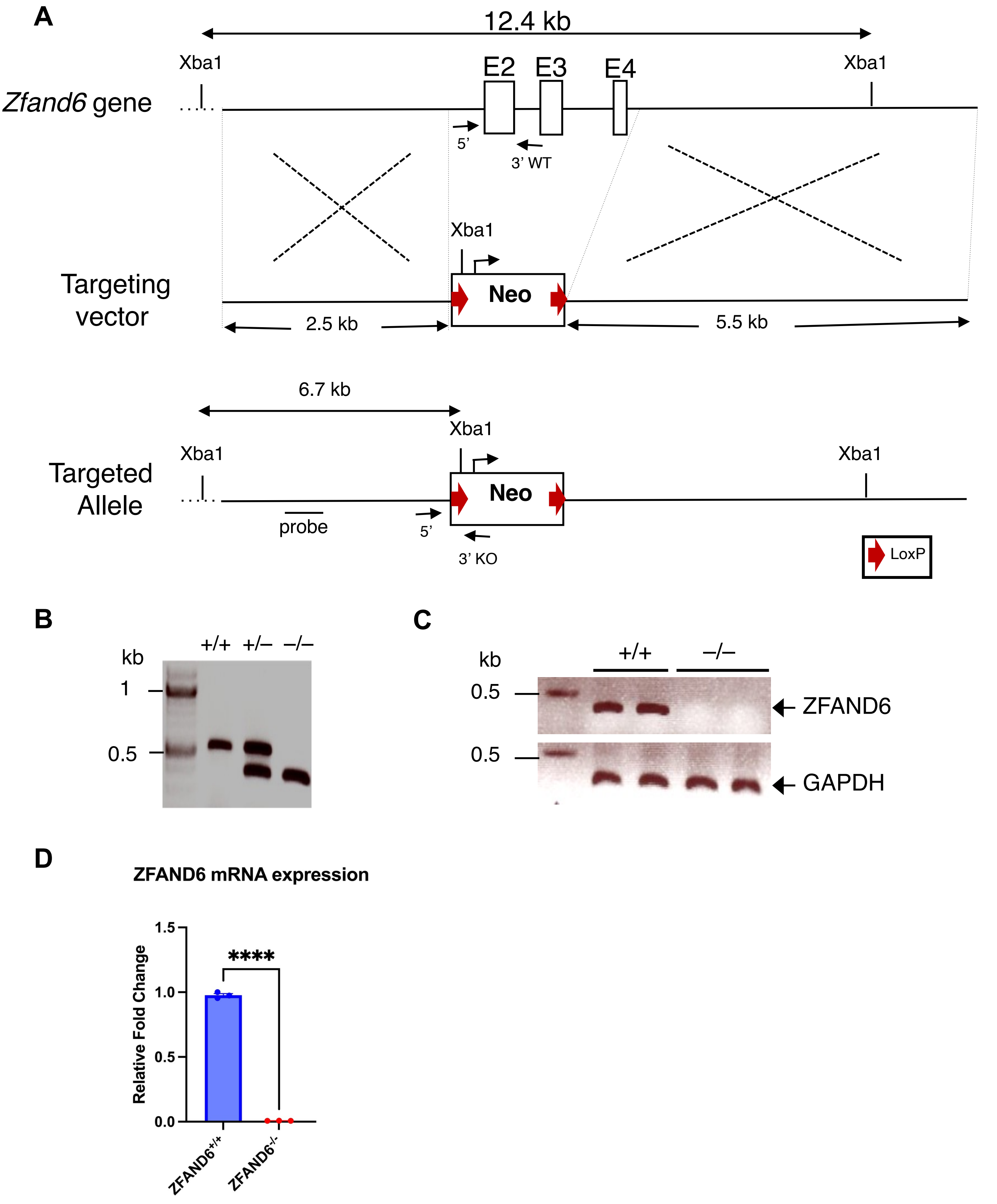

**Extended Data Fig. 1. Generation of *Zfand6*$^{–/–}$ mice, MEFs and BMDMs. (A)** Schematic of mouse *Zfand6* genomic organization, targeting vector and targeted allelle after homologous recombination to remove exons 2-4. **(B)** Genotyping of mouse tail DNA using the primers indicated in panel A. **(C)** RT-PCR of ZFAND6 and GAPDH amplicons using RNA extracted from *Zfand6*$^{+/+}$ and *Zfand6*$^{–/–}$ MEFs. **(D)** qRT-PCR of *Zfand6* in WT and *Zfand6*$^{–/–}$ BMDMs. Data presented as mean ± SEM. Unpaired Student's t-test. ****P<0.0001.

### Extended Data Fig. 2

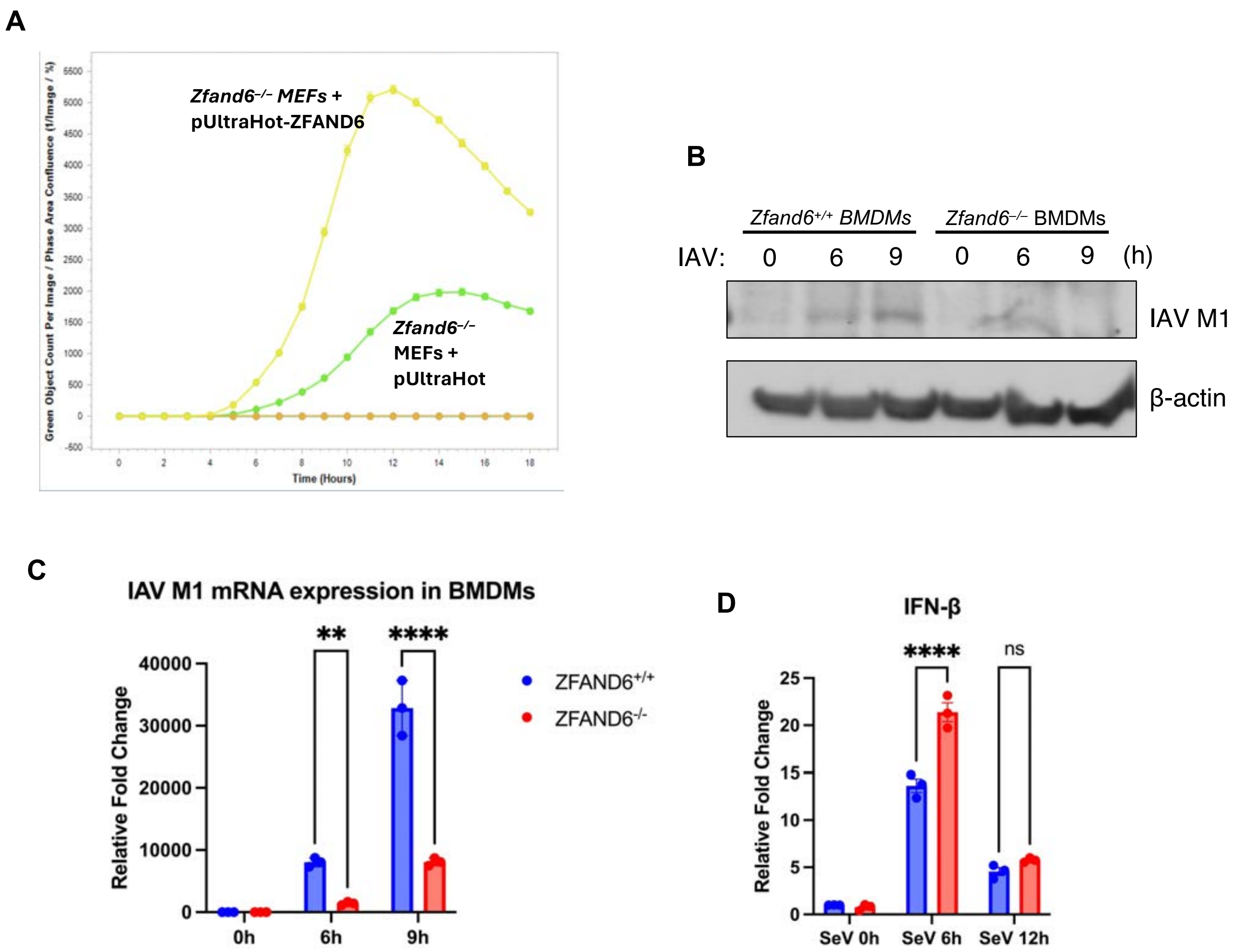

**Extended Data Fig. 2. The absence of ZFAND6 promotes resistance to virus infection. (A)** Incucyte S3 live-cell imaging of *Zfand6*$^{-/-}$ MEFs expressing pUltra-Hot empty vector or pUltra-Hot-ZFAND6 infected with VSV-GFP (MOI=1). **(B)** WT and *Zfand6*$^{-/-}$ BMDMs were infected with IAV (MOI=1) for 0, 6, or 9 hr. Protein lysates were subjected to western blotting for M1 and b-actin. **(C)** qRT-PCR of M1 in WT and *Zfand6*$^{-/-}$ BMDMs infected with IAV (MOI=1). **(D)** qRT-PCR of IFN-β in WT and *Zfand6*$^{-/-}$ MEFs infected with SeV (20 HA/mL) at the indicated time points. Data presented as mean ± SEM. Unpaired Student's t-test. **P<0.01, ****P<0.0001, ns=not significant.

### Extended Data Fig. 3

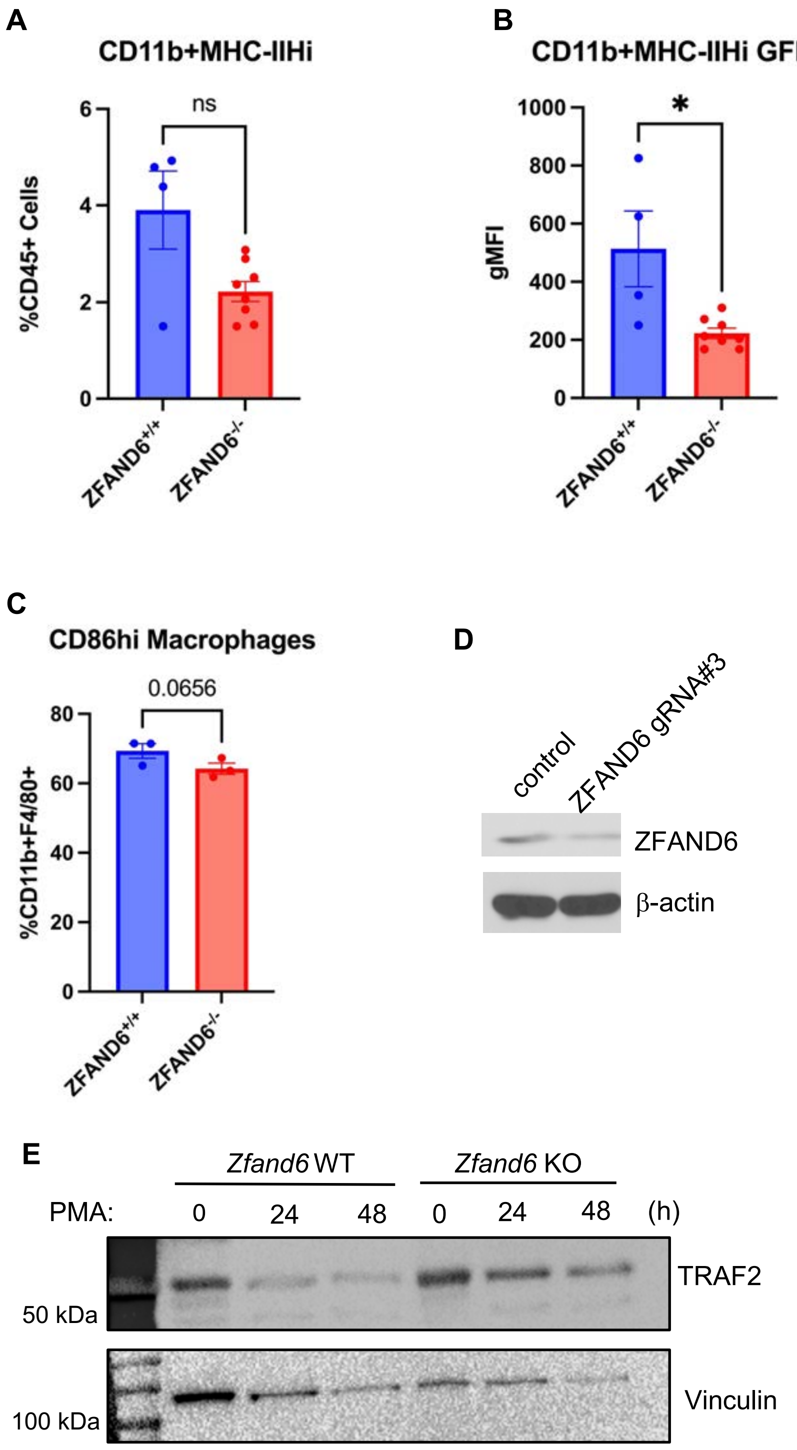

**Extended Data Fig. 3. ZFAND6 deficiency impairs the expression of myeloid cell activation markers and TRAF2 degradation.** *Zfand6*$^{+/+}$ x Mx1-GFP and *Zfand6*$^{-/-}$ x Mx1-GFP mice were infected with 1000 FFC of IAV (PR8). Flow cytometry was performed to detect CD11b+MHC-II$^{hi}$ cells **(A)**, CD11b+MHC-II$^{hi}$GFP+ cells **(B)** and GFP fluorescence intensity in CD11b+MHC-II$^{hi}$ cells. **(C)** Population of CD86$^{hi}$ cells from CD11b+F4/80+ BMDMs. **(D)** Western blot with the indicated antibodies using lysates from the bulk population of THP-1 cells expressing control gRNA or ZFAND6 gRNA3. **(E)** WT or THP-1 ZFAND6 KO cells were treated with PMA for the indicated times and lysates were subjected to western blotting with anti-TRAF2 and anti-Vinculin. Data presented as mean ± SEM. Unpaired Student's t-test. *P<0.05, ns=not significant.

### Extended Data Fig. 4

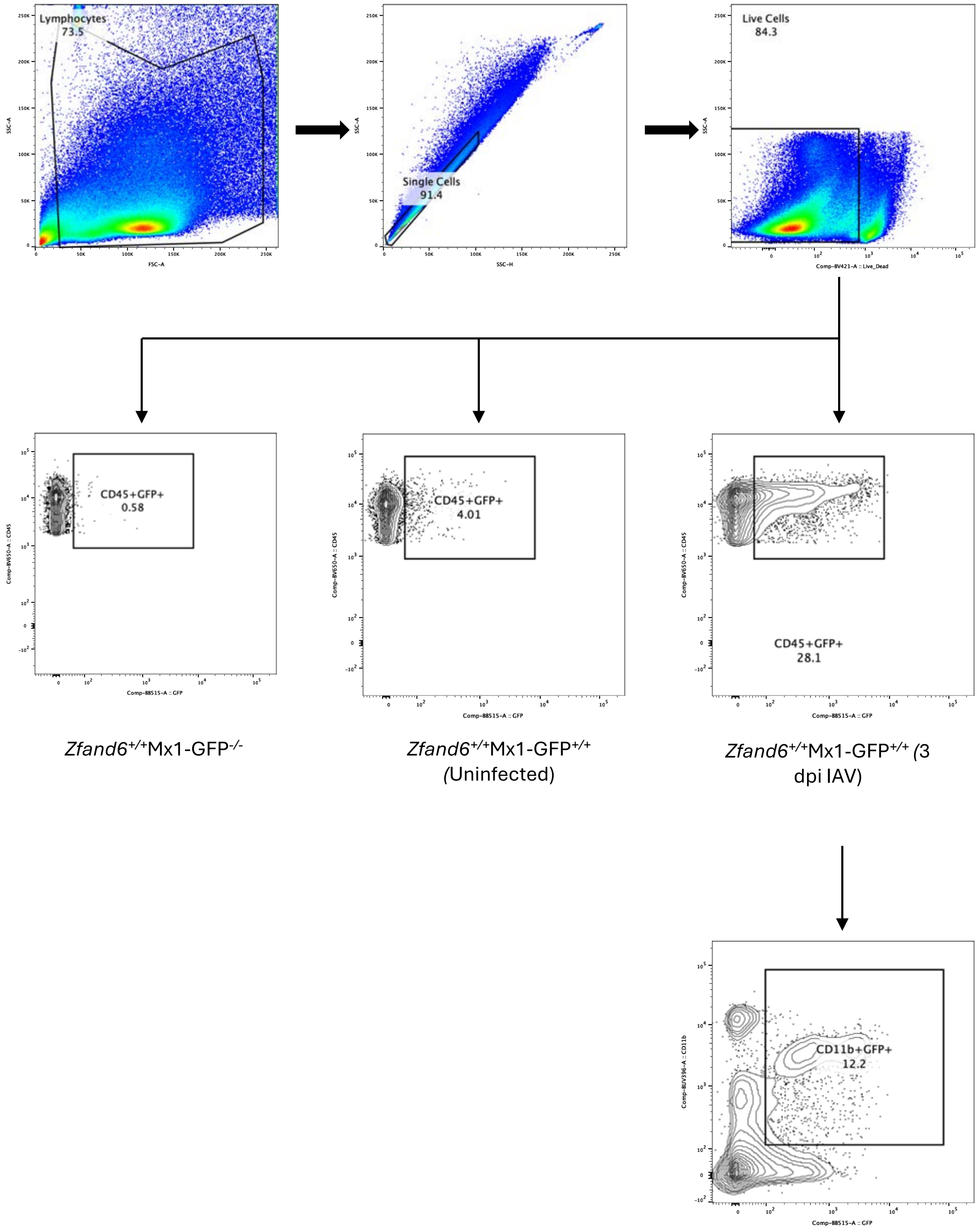

**Extended Data Fig. 4. Gating strategy for flow cytometric analysis of lung homogenates.** Gating strategy for single cell lung homogenates from IAV-infected and uninfected *Zfand6*$^{+/+}$ x Mx1-GFP, *Zfand6*$^{–/–}$ x Mx1-GFP and *Zfand6*$^{+/+}$ mice for flow cytometric analysis.

### Extended Data Fig. 5

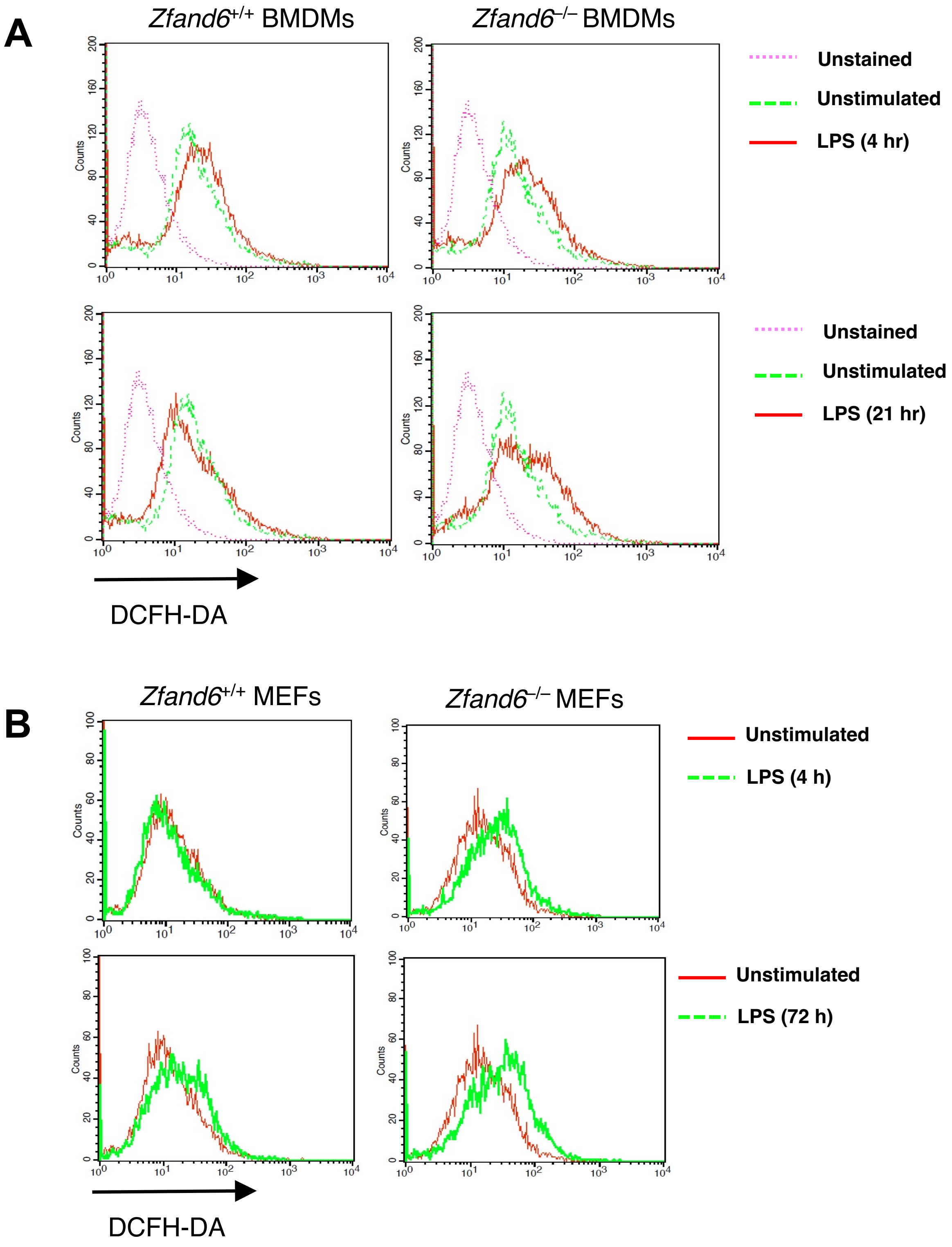

**Extended Data Fig. 5. ZFAND6 deficiency promotes increased LPS-induced ROS production. (A)** Flow cytometry of *Zfand6*$^{+/+}$ and *Zfand6*$^{–/–}$ BMDMs stimulated with LPS (20 ng/ml) for the indicated times and stained with DCFH-DA. **(B)** Flow cytometry of *Zfand6*$^{+/+}$ and *Zfand6*$^{–/–}$ MEFs stimulated with LPS (300 ng/ml) for the indicated times and stained with DCFH-DA.

### Extended Data Fig. 6

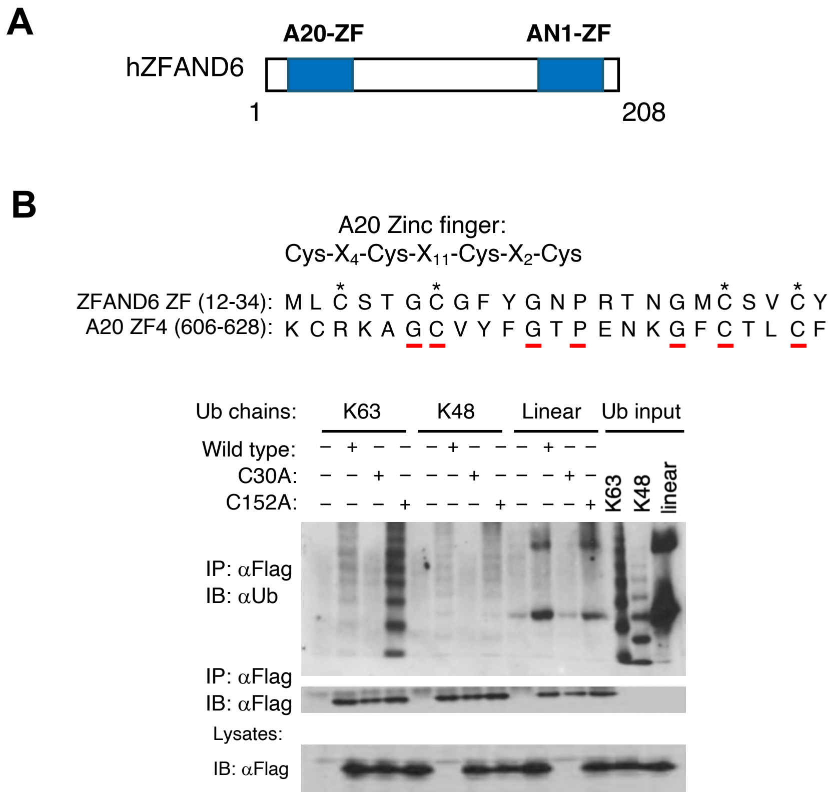


**Extended Data Fig. 6. ZFAND6 interacts with polyubiquitin chains. (A)** Schematic of ZFAND6 domains. **(B)** Sequence alignment of ZFAND6 A20-like ZF domain and A20 ZF4. Co-IP of lysates from 293T cells transfected with Myc-Flag-ZFAND6, ZFAND6 C30A and ZFAND6 C152A incubated with recombinant K63, K48 or linear poly-Ub chains. IP was performed with anti-Flag and IB with anti-Ub.
